# microRNA-1 represses signaling pathway components to impact embryonic structures derived from all three germ layers

**DOI:** 10.1101/2022.02.27.482171

**Authors:** Nina Faye Sampilo, Jia L. Song

## Abstract

microRNAs are evolutionarily conserved non-coding RNAs that direct post-transcriptional regulation of target transcripts. We use the sea urchin embryo to achieve a comprehensive understanding of miR-1’s function in a developing embryo. Results indicate that miR-1 regulates gut contractions, specification, and positioning of serotonergic neurons, as well as mesodermally-derived muscles, pigment cells, and skeletogenic cells. Gain-of-function of miR-1 generally leads to more severe developmental defects than its loss-of-function. We identified that miR-1 directly suppresses *Ets1/2, Tbr,* and *VegfR7* of the skeletogenic gene regulatory network, and *Notch, Nodal,* and *Wnt1* signaling components. We found that miR-1’s direct suppression of *Nodal* may indirectly regulate *FoxQ2* to impact serotonergic neurons. Excess miR-1 may lead to decreased *Nodal* and *Notch* that result in decreased circumpharnygeal muscle fibers and the number of pigment cells. The striking ectopic skeletal branching induced by miR-1 mimic injections may be explained by miR-1’s direct suppression of *Nodal* that leads to expression changes of *Vegf3,* and *Fgfa* that mediate skeletogenesis. This work demonstrates that miR-1 plays a diverse regulatory role that impacts tissues derived from all germ layers.

**Summary statement:** This study identifies wide-ranging regulatory roles and regulatory mechanisms of miR-1 that impact structures derived from all three germ layers during embryonic development.

## Introduction

microRNA-1 (miR-1) is classified as a myomiR because of its enriched expression in vertebrate muscle tissue and its function in myogenesis, angiogenesis, and vascularization (Mansfield et al., 2004; Sokol and Ambros, 2005; Wienholds et al., 2005; Zhao et al., 2005; McCarthy, 2011). miR-1 has also been implicated in regulating neuronal growth and survival by directly targeting *Bdnf, Hes1*, and potentially *Hsp-70* (Varendi et al., 2014; Chang et al., 2016; Yi et al., 2016; Zheng et al., 2017). While there are extensive studies demonstrating miR-1’s role in angiogenesis and myogenesis, the exact molecular regulatory mechanisms of miR-1 are still not fully understood. Additionally, a systematic analysis of both loss and gain-of-function of miR-1 in a developing embryo is still lacking.

The miR-1 sequence is highly conserved amongst metazoans (Fig. S1). Most mammalian genomes contain two copies of miR-1 that perform similar functions (Heidersbach et al., 2013). Previously, we found that miR-1 is one of the most expressed miRNAs in the purple sea urchin embryo (Song et al., 2012). The sea urchin has ∼50 annotated miRNAs, which is relatively small in contrast to the 519 miRNAs in humans (Song et al., 2012; Fromm et al., 2015). The sea urchin embryo contains a single miR-1, making it feasible to use this embryo to provide a deeper understanding of miR-1’s function in development.

The developmental processes of the sea urchin and humans are remarkably similar at the cellular and molecular level (Adonin et al., 2020). Similar to vertebrates, they utilize highly conserved signaling pathways such as the canonical Wnt (cWnt)/β-catenin signaling for anterior-posterior (AP) axis formation (Wikramanayake et al., 1998; Angerer and Angerer, 2000; Komiya and Habas, 2008) and Nodal/Bmp signaling for specification of the dorsal-ventral (DV) secondary body axis (Duboc et al., 2004; Furtado et al., 2008; Chhabra et al., 2019). Immediately after fertilization, maternal inputs, zygotic transcription, and signaling mechanisms help define distinct gene regulatory networks (GRNs) (Fig. 1A) (Logan et al., 1999; Martik et al., 2016). By the mesenchyme blastula stage (24 hours post fertilization; hpf), germ layer specification has already occurred;during gastrulation, the three germ layers are differentiated by cross-regulation between signaling pathways, GRN interactions (Lyons et al., 2012), as well as post-transcriptional regulation mediated by miRNAs (Stepicheva and Song, 2015; Sampilo et al., 2018; Sampilo et al., 2021).

**Figure 1.**
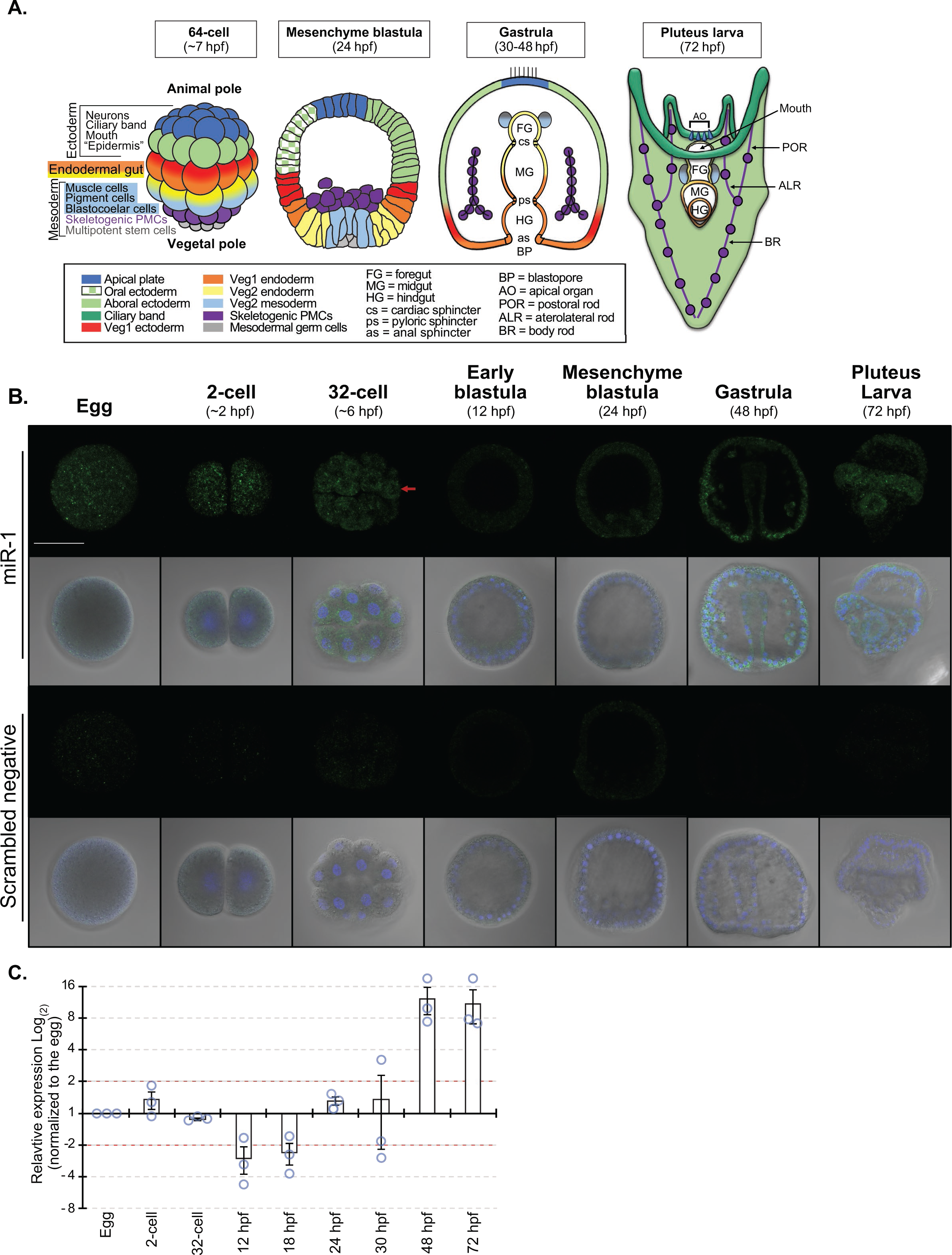
miR-1 is dynamically expressed throughout development. (A) Schematics depicting various cell fates in various developmental stages of the sea urchin embryo. (B) F*IS*H was used to detect miR-1 at various developmental stages and counterstained with DAPI against DNA (blue). miR-1 is expressed perinuclearly in the 32-cell stage embryo (arrow). miR-1 has increased expression during gastrula stage and is enriched in the larval ciliary band and gut. Larva: N=13, 3 biological replicates. Scale bar = 50 μm. (C) miR-1 expression is measured by relative miRNA RT-qPCR at various developmental stages. Red dashed lines indicate 2-fold expression difference. Blue circles represent datum points. 3 biological replicates.

Although the body plans and structures of deuterostomes are diverse, the sea urchin embryo and vertebrates utilize conserved factors for analogous structures. For example, the sea urchin skeletogenesis is thought to be analogous to vertebrate angiogenesis and vascularization since these processes utilize a common set of transcription factors (TFs) (Ets1/2, Erg, Hex, Tel, and FoxO) and signaling pathways (Vegf, Nodal/Bmp, Notch, and Angiopoetin) (Duboc et al., 2005; Morgulis et al., 2019). In response to the Vegf ligand in the ectoderm, the sea urchin *VegfR*-expressing skeletogenic primary mesenchyme cells (PMCs) initiate the formation of the skeletal rudiment, differentiate, and pattern (Ettensohn and McClay, 1986; Duloquin et al., 2007; McIntyre et al., 2014). The migrating PMCs form a syncytium, connected by filopodial membranes between cell bodies where biomineralization enzymes form calcite granules (Ettensohn and McClay, 1986; Armstrong and McClay, 1994; Wilt, 2002). Similarly, the sea urchin larva utilizes similar TFs as vertebrates (FoxA, GataE, Xlox, Cdx) critical for gut differentiation (Annunziata et al., 2019). The tripartite gut is compartmentalized with the cardiac, pyloric, and anal sphincters (Fig. 2B) (Yaguchi and Yaguchi, 2019). For the nervous system, the sea urchin embryo has serotonergic neurons and ganglia located in the apical organ that is analogous to the vertebrate central nervous system and sensory and motor neurons within the ciliary band that is analogous to the peripheral nervous system that control swimming and feeding behaviors (Yaguchi and Katow, 2003; Wood et al., 2018; Yaguchi and Yaguchi, 2021). Additionally, the sea urchin embryo uses orthologous neuronal transcriptional factors as those expressed in the vertebrate forebrain (Six3, ZIC2, Achaete-scute, NKX2.1 and FEZ) (Wei et al., 2009). Thus, using the sea urchin as a simple and experimentally tractable organism, we can better understand complex molecular mechanisms.

**Figure 2.**
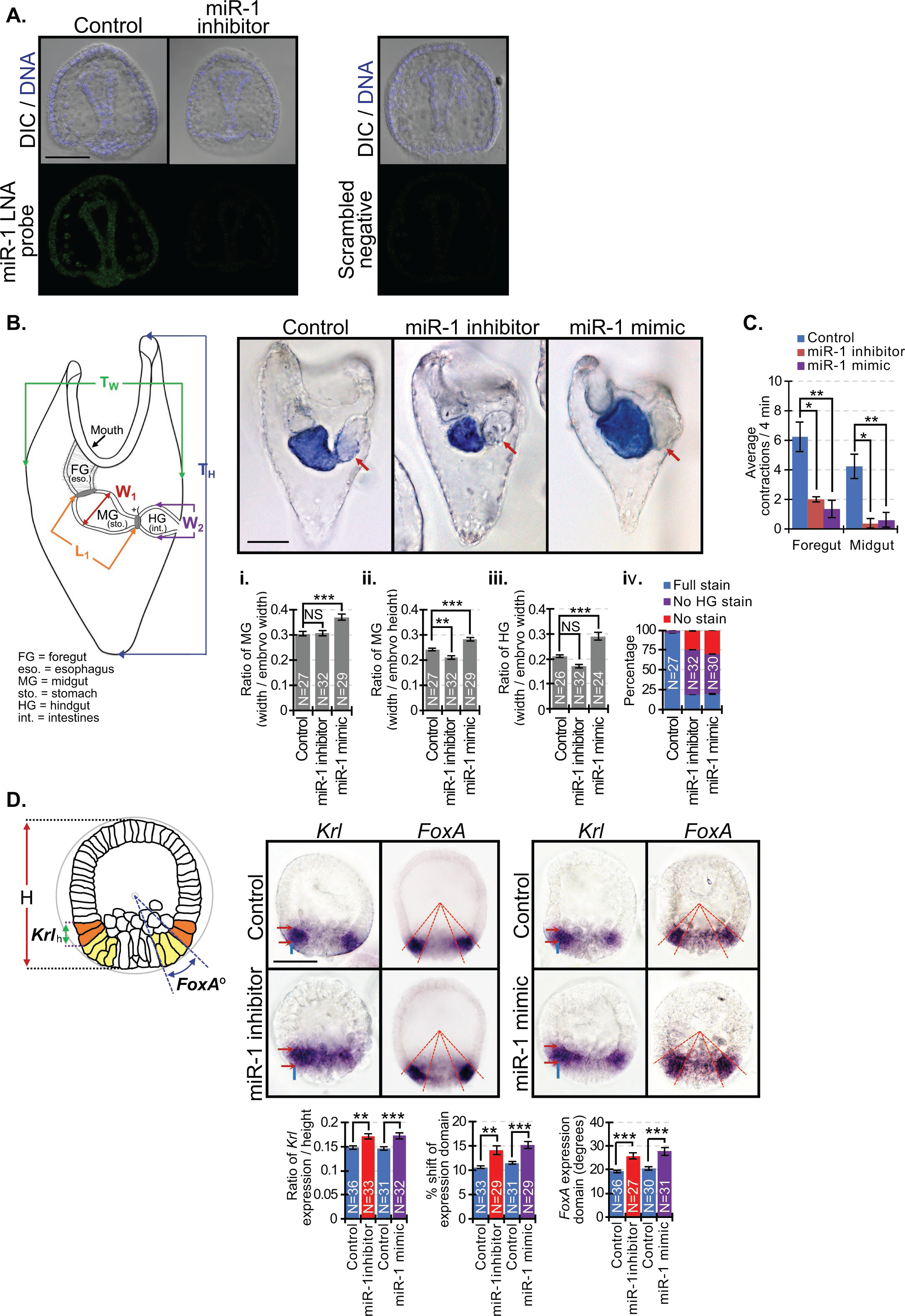
Perturbation of miR-1 results in gut defects. (A) miR-1 inhibition resulted in undetectable level of miR-1 compared to the control. N=20 (Bi-iii) Width of MG and width of HG of larvae were measured (W_1_ and W_2_), and their ratios to overall width of larvae (T_w_) were calculated. The length of MG was measured (L_1_) and calculated as the ratio to overall height of larvae (T_H_). Compared to control, miR-1 inhibition resulted in shorter MGs while miR-1 mimic-injected larvae have much wider and longer MGs and HGs. (Biv) Control and miR-1 perturbed larvae were assessed for alkaline phosphatase activity (blue stain). (C) miR-1 perturbed larvae exhibit significantly less contractions of the foregut and the midgut compared to control. 2 biological replicates. (D) Blastulae were hybridized with *Krl* and *FoxA* RNA *in situ* probes. *Krl* is expressed in the Veg1 endoderm (orange cells in schematic). The spatial expression domains (red arrows) and anterior shift (blue lines) of *Krl* were measured on both sides of each embryo. *FoxA* is expressed in the Veg2 endoderm (yellow cells in schematic), which are positioned at an angle (red dashed lines). Student’s t-test. 2 biological replicates. Scale bar = 50 µm.

We discovered that miR-1 regulates the endodermally-derived gut contractions, specification and positioning of ectodermally-derived serotonergic neurons, as well as mesodermally-derived circumpharyneal muscles, pigment cells (PCs), and skeletogenesis. Interestingly, gain-of-function of miR-1 leads to more severe developmental defects than its loss-of-function. We further identified miR-1 to directly suppress *Ets1/2, Tbr,* and *VegfR7* of the PMC GRN, and signaling components *Notch, Nodal,* and *Wnt1.* We found that miR-1’s direct suppression of *Nodal* may indirectly regulate *FoxQ2* to impact serotonergic neurons. miR-1 mimic-induced overexpression may lead to decreased *Nodal* and *Notch* that result in decreased circumpharnygeal muscle fibers and the number of pigment cells. Additionally, excess miR-1 also resulted in numerous ectopic skeletal branching that may be explained by miR-1’s direct suppression of *Nodal* that leads to expression changes of *Vegf3* and *Fgfa* that mediate patterning and skeletogenesis, respectively. This study identifies wide-ranging regulatory roles and regulatory mechanisms of miR-1 that impact structures derived from all three germ layers during embryonic development.

## Experimental Procedures

### Animals

Adult purple sea urchin, *Strongylocentrotus purpuratus* (*Sp*), were obtained from Point Loma Marine Invertebrate Lab, (Lakeside, CA) and Marinus Scientific, LLC (Long Beach, CA). Adult males and females were intracoelomically injected with 0.5 M KCl to obtain sperm and eggs. Filtered natural seawater (FSW) (collected from Indian River Inlet; University of Delaware) or artificial seawater (ASW) made from Instant Ocean© was used for embryo cultures incubated at 15°C.

### Cloning

For generating 3’UTR luciferase reporter constructs of predicted miR-1 targets, PCR primers, or FragmentGENE DNA fragments (Genewiz, South Plainfield, NJ) were designed based on sequence information available from the sea urchin genome (echinobase.org) (Table S1). Amplified PCR products of *Bmp2/4, Ets1/2, Notch, Nodal,* and *Tbr* 3’UTRs were first cloned into ZeroBlunt vector (Thermo Fisher Scientific, Waltham, MA), and then subcloned into the *Renilla* luciferase (*R*luc) reporter construct. Wildtype (WT) constructs of *IgTM* and *Nodal* were commercially synthesized prior to cloning. *VegfR7* and *Wnt1* 3’UTRs were previously cloned (Stepicheva and Song, 2015; Sampilo et al., 2021). Mutations were generated within the miR-1 seed sequences using the QuikChange Lightning or QuikChange Multi Site-Directed Mutagenesis Kit (Agilent Technologies, Santa Clara, California) to disrupt miR-1’s binding and regulatory function (Gregory et al., 2008; Stepicheva et al., 2015). The predicted miR-1 seed sites (canonical site: 5’ ACATTCC 3’) within the 3’UTRs of *Ets1/2, Dri, Tbr, VegfR7, Notch, Nodal, Bmp2/4, Wnt1,* and *IgTM* were modified at the third and fifth base pair (Table S1). We have previously demonstrated that truncated miRNA seed sequence differing in one nucleotide at the 5’ end is sufficient in miRNA-mRNA target recognition and function (Sampilo et al., 2021). *Nodal* and *Wnt1* contain truncated miR-1 sites (6 out of 7 bps), and *Bmp2/4* and *IgTM* contain mismatched miR-1 seed sites which differ in one nucleotide within the seed sequence (Table. S1). Only two of the three potential miR-1 binding sites within *Nodal* 3’UTR were mutated due to sequence complexity. Firefly luciferase was used as a normalization control as previously described (Stepicheva et al., 2015). Each of the construct sequences were verified by DNA sequencing (Genewiz, South Plainfield, NJ). Luciferase constructs containing *Ets1/2, VegfR7, Nodal,* and *IgTM* 3’UTRs were linearized with *EcoR*I, while *Tbr, Bmp2/4, and Wnt1* 3’UTRs were linearized with *Not*I. The luciferase constructs and *Firefly* luciferase mRNA were *in vitro* transcribed using the Sp6 mMessage machine kit (Ambion Inc, Austin, Texas). *In vitro* transcribed mRNAs were purified using Macherey-Nagel Nucleospin® RNA Clean-up kit (Macherey-Nagel, Bethlehem, PA) prior to injections.

### Microinjections

Microinjections were performed as previously described (Cheers and Ettensohn, 2004; Stepicheva and Song, 2014) with modifications. All injection solutions were prepared in a 2.5 µl solution consisting of 20% glycerol and 0.4 μg/µl of 10,000 MW neutral non-fixable Texas Red dextran (Thermo Fisher Scientific, Waltham, MA). Approximately 1-2 picoliter was injected into each newly fertilized egg based on the size of the injection bolus at about one-fifth of the egg diameter. miR-1 miRCURY LNA miRNA Power Inhibitor and miRCURY LNA miRNA mimic were obtained from Qiagen (Germantown, MD). miR-1 inhibitor (Hsa-miR-1-3p, ID# YI04100840; 5’-UGGAAUGUAAAGAAGUAUGUAU-3’) and miR-1 miRCURY LNA miRNA mimic (ID#YM00472818; 5’-UGGAAUGUAAAGAAGUAUGUAU-3’) were used at 10 µM, 30 µM and 40 µM concentrations. Cel-miR-39-3p LNA mimic (ID#YM00479902; 5’-UCACCGGGUGUAAAUCAGCUUG-3’) was used as an injection control (not present in the sea urchin or humans). miR-1 inhibitor and miR-1 mimic were co-injected at a 1:1 molar ratio (40 µM miR-1 inhibitor + 40 µM miR-1 mimic) to test the specificity of miR-1 inhibitor.

### Dual-luciferase quantification

The injection solutions for the dual-luciferase assay contained 20% sterile glycerol, 0.4 μg/µl 10,000 MW Texas Red lysine-charged dextran, 100-200 ng/μl *Firefly* mRNA, and 100 ng/μl *R*luc mRNA (*Ets1/2, Dri, Tbr, VegfR7, Notch, Nodal, Bmp2/4, Wnt1,* and *IgTM*). 20-50 embryos were collected at the mesenchyme blastula stage (24 hpf). Dual-luciferase assays were performed using the Promega Dual-Luciferase Reporter (DLR™) Assay Systems with the Promega GloMax 20/20 Luminometry System (Promega, Madision, WI) (Stepicheva et al., 2015; Stepicheva and Song, 2015; Sampilo et al., 2018; Sampilo et al., 2021). The *R*luc values were normalized to the Firefly signal to account for microinjection volume differences. *R*luc data with mutated miR-1 seed sites were normalized to the *R*luc with wildtype (WT) 3’UTR constructs. P-value was analyzed using Student’s t-test. All error bars represent Standard Error (SEM).

### Immunofluorescence

Gastrulae and larvae were fixed in 4% paraformaldehyde (PFA) (20% stock; EMS, Hatfield, PA) in FSW overnight at 4°C (1D5, Serotonin, and Sp1), or fixed with 3.7% PFA for 20 min at RT followed by 1 min post-fix with ice-cold methanol (for SynB). 10X Phosphate Buffered Saline (PBS) (Bio-Rad, Hercules, CA) was diluted at 1:10. Three PBS-Tween (0.05% Tween-20 in 1X PBS for 1D5) or PBS-Triton (0.1% Triton in 1X PBS for Serotonin, Sp1, and SynB) washes were performed, followed by 1 h block with 4% sheep serum (MilliporeSigma, St. Louis, MO). 1D5 antibody was used at 1:50 to visualize PMCs (McClay et al., 1983); serotonin at 1:500 (MilliporeSigma, Cat#S5545) to visualize serotonergic neurons; Sp1 at 1:200 to visualize pigment cells (Developmental Studies Hybridoma Bank, Iowa City, Iowa, Cat#531884); and SynB at 1:200 to visualize mature and functional neurons (Burke et al., 2006; Leguia et al., 2006) (gift from Dr. Gary Wessel, Brown University). Antibodies were diluted in PBS-Tween/Triton with 4% sheep serum, and embryos were incubated overnight to 3 days at 4°C and washed 3 times with PBS-Tween/Triton, followed by goat-anti mouse (1D5 and Sp1) or anti-rabbit (Serotonin and SynB) secondary antibody (Thermo Fisher Scientific, Waltham, MA) at 1:300 for 1 h at RT. Embryos were then washed 3 times with PBS-Tween/Triton and mounted on slides for confocal imaging. For visualization of DNA, embryos were counterstained with Hoechst dye (Lonza, Walkersville, MD), DAPI (Thermo Fisher Scientific, Waltham, MA), or VECTASHIELD® Antifade Mounting Medium with DAPI (Vector Laboratories, Burlingame, CA).

### Whole mount *in situ* hybridization (WM*IS*H)

Partial coding sequences of *Bmp2/4, FoxA, FoxQ2, Krl, Nodal, Not1, Vegf3,* and *Wnt1* were cloned into ZeroBlunt vector to generate RNA probes (Thermo Fisher Scientific, Waltham, MA) Constructs were linearized using FastDigest™ (Thermo Fisher Scientific, Waltham, MA) and *in vitro* transcribed with DIG RNA Labeling Kit (Millipore Sigma, St. Louis, MO) (Table S2). *FoxA, Krl, Vegf3* and *Wnt1* RNA probes were previously cloned (Stepicheva and Song, 2015; Sampilo et al., 2021). WMISH was conducted according to previous publications (Arenas-Mena et al., 2000; Minokawa et al., 2004; Stepicheva et al., 2015; Stepicheva and Song, 2015). Probes were used at 1 ng/µl and incubated at 50°C for 5-7 days.

To test endodermal differentiation, we examined the activity of alkaline phosphatase (Kumano and Nishida, 1998; Drawbridge, 2003). Larvae 5 days post fertilization (5 dpf) were fixed in MOPS-paraformaldehyde based fixative (4% paraformaldehyde, 100 mM MOPS pH 7, 2 mM MgSO_4_, 1 mM EGTA, and 0.8 M NaCl) for 10 min at RT as previously described (Stepicheva et al., 2015). Embryos were then washed with alkaline phosphatase buffer 3 times (100 mM Tris pH 9.5, 100 mM NaCl, 50 mM MgCl_2_, 0.1% Tween-20), followed by staining until the desired color was developed with the staining solution (0.1 M Tris pH 9.5, 50 mM MgCl_2_, 0.1 M NaCl, 1 mM Levamisole, 10% Dimethylformamide, 50 mg/ml NBT in 70% dimethyl formamide and 50 mg/ml BCIP in 100% dimethyl formamide). Staining was stopped with multiple washes of MOPS buffer (0.1 M MOPS pH 7, 0.5M NaCl, and 0.1% Tween-20).

### Imaging and Phenotyping

Representative images were taken with Zeiss LSM 880 scanning confocal microscope using Zen software or ZEISS Observer Z1 using AxioVision software (Carl Zeiss Microscopy, LLC, White Plains, NY). For videos of pigment cells and gut contractions, live embryos were collected 5 dpf and mounted in FSW onto protamine sulfate (PS)-coated coverslips, creating a positively-charged surface (Stepicheva and Song, 2014). For live behavior examination, control injected, miR-1 inhibitor, and miR-1 mimic-injected embryos were mounted on the same multichambered PS-coated coverslip to avoid variability of environmental conditions and response. To measure dorsoventral connecting rod (DVC) length or PMC migration, ZEISS Observer Z1 microscope was used to take Z-stacks of differential interference contrast (DIC) and 1D5-immunolabeled images. ZEISS AxioCam105 color camera was used to take *in situ* images. AxioVision or Zen 3.1 software (Carl Zeiss Microscopy, White Plains, NY) was used to measure the length of DVCs, PMC migration distance, *in situ* expression domains and to determine the center of gastrulae in vegetal views to measure angles of *Vegf3, Wnt1, Nodal, Not1,* and *Bmp2/4* expression domains, ventral ectodermal (VE), and dorsal ectodermal (DE) domains. N is the total number of embryos examined unless otherwise stated. NS = not significant, *p < 0.05, **p < 0.001, ***p < 0.0001. All error bars represent SEM.

To measure the average intensity levels of arbitrary fluorescent units (intensity) in larvae immunolabeled with serotonin and SynB, maximum intensity projection images were analyzed with ImageJ (Schneider, 2012; Remsburg et al., 2021). Serotonin and SynB expression within the ciliary band was measured and the background was subtracted.

### Real-Time, quantitative PCR (qPCR)

To examine the levels of endogenous miR-1 expression within a developing embryo, 200-500 embryos were collected at various developmental stages (Fig. 1C). Purification of total RNA was done using miRNAeasy Micro Kit (QIAGEN, Germantown, MD). cDNA synthesis of 100 ng total RNA was performed with miRCURY LNA RT Kit (10 µl volume reaction) which adds a 5’ universal tag of a poly(A) tail to mature miRNA templates (QIAGEN, Germantown, MD). cDNA template was diluted 1:10, and miRNA qPCR was performed using miRCURY LNA miRNA PCR Assays (QIAGEN, Germantown, MD) in QuantStudio 6 Real-Time PCR cycler system (Thermo Fisher Scientific, Waltham, MA). Sea urchin miR-200 were used as normalization controls due to its similar expression from the cleavage to the larval stages (Song et al., 2012). Results are shown as fold changes compared to the egg stage using the Ct^-2ΔΔ^ method as previously described (Stepicheva et al., 2015). miRCURY LNA miRNA PCR Primer Mix is against human miR-1 (Hsa-miR-1-3p).

To measure the transcriptional changes of key TFs for ectodermal, endodermal, and mesodermal specification, transcripts that encode TFs of the skeletogenic GRN, and biomineralization enzymes, we injected zygotes with control, miR-1 inhibitor, and miR-1 mimic. 100 of these blastulae were collected at 24 hpf. Total RNA was extracted by using the Macherey-Nagel Nucleospin® RNA Clean-up XS kit (Macherey-Nagel, Bethlehem, PA). cDNA was synthesized using iScript cDNA synthesis kit (Bio-Rad, Hercules, CA). qPCR was performed using 2.5 embryo equivalents for each reaction with the Fast SYBR or PowerUp Green PCR Master Mix (Thermo Fisher Scientific, Waltham, MA) in the QuantStudio 6 Real-Time PCR cycler system (Thermo Fisher Scientific, Waltham, MA). Results were normalized to the mRNA expression of ubiquitin and depicted as fold changes compared to control embryos using the Ct^-2ΔΔ^ method as previously described to analyze the relative changes in gene expression (Stepicheva et al., 2015). Primer sequences were designed using the Primer 3 Program (Rozen and Skaletsky, 2000) and are listed in Table S3. 3-6 biological replicates were conducted. Statistical significance was calculated using two-tailed unpaired Student’s t-tests.

## Results

### Expression of miR-1 peaks at gastrula stage and is enriched in the larval gut and ciliary band

miR-1 is expressed maternally. At the 32-cell stage, the expression of miR-1 is enriched in the perinuclear region (Fig. 1B). miR-1 expression peaks at the gastrula stage with ubiquitous expression. At the larval stage, miR-1 is enriched in the gut and ciliary band (Fig. 1B). These results are consistent with relative quantitative levels of miR-1 using real-time, qPCR (Fig. 1C).

### miR-1 modulates endodermal differentiation and gut function

To examine the function of miR-1 in early development, we injected miR-1 inhibitor for loss-of-function studies or miR-1 mimic for gain-of-function studies. To test the efficacy of the miR-1 inhibitor, zygotes were injected with control or miR-1 inhibitor, followed by miR-1 F*I*SH (Fig. 2A). We observed a reduction of miR-1 in 100% of miR-1 inhibitor-injected gastrulae, indicating the effectiveness of the miR-1 inhibitor.

Results indicate that miR-1 perturbed embryos have aberrant gut morphology compared to the control (Fig. 2B). In miR-1 inhibitor-injected larvae, width of midgut (MG) is smaller compared to control (Fig. 2Bi). Conversely, in miR-1 mimic-injected larvae, width of MG and hindgut (HG), in addition to length of MG is much wider compared to control (Fig. 2Bi-iii). We further used alkaline phosphatase (ALP) staining as a marker for endodermal differentiation (Drawbridge, 2003; Stepicheva et al., 2015). Control-injected embryos have ALP activity within the MG and part of the HG, whereas miR-1 perturbed larvae exhibited a loss of ALP within the HG and/or in the MG (Fig. 2Biv). Interestingly, both miR-1 inhibitor and miR-1 mimic-injected larvae have significantly less foregut and hindgut contractions compared to the control, indicating a potential defect in gut functions (Fig. 2C, Movies 1-3).

We examined the spatial expression of *Krl* and *FoxA* that are important for specifying the foregut and midgut (Oliveri et al., 2006; Yamazaki et al., 2008). In miR-1 perturbed embryos, the spatial expression of *Krl* and *FoxA* expanded, and *Krl*’s expression also shifted anteriorly compared to control (Fig. 2D). The spatial expression changes of key TFs involved in endodermal specification could in part contribute to the morphological changes in the larval gut. Results indicate that miR-1 may regulate gut morphology, differentiation, and function.

### Perturbation of miR-1 results in circumpharyngeal muscle defects

Since defects in gut contraction may attribute to gut muscle defects, we examined the structure of the circumpharyngeal muscles (Fig. 3). The number of filamentous actin-rich fibers (F-actin) within the muscle fiber ring was significantly decreased and less structured in both miR-1 perturbed larvae compared to the control. Almost 20% of miR-1 mimic-injected larvae had a complete loss of detectable F-actin (Fig. 3). The miR-1 inhibitor-induced defects were rescued by co-injection with the miR-1 mimic, indicating that the muscle fiber defects are due to miR-1 perturbation.

**Figure 3.**
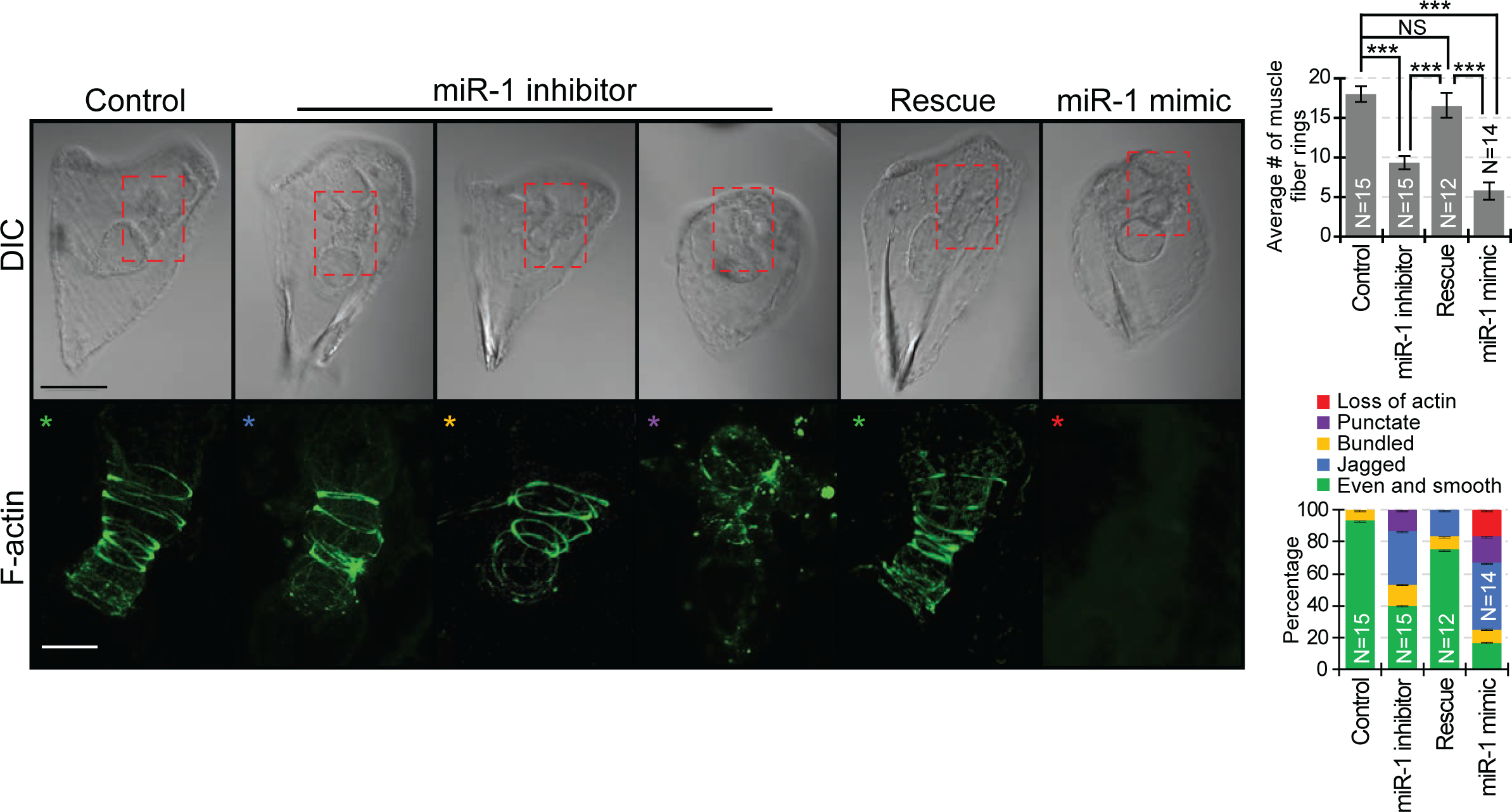
Perturbation of miR-1 results in circumpharyngeal muscle morphological defects. Circumpharyngeal muscles were labeled with Phalloidin to detect F-actin. Z-stack of confocal images were taken to count muscle fiber rings. Compared to control, miR-1 perturbed larvae exhibited less or a complete loss of F-actin, in addition to irregular F-actin morphology (boxed areas). miR-1 inhibitor induced defects were rescued by co-injection of miR-1 mimic. Colored asterisks correspond to phenotypes in the bar graph. 4 biological replicates. Student’s t-test. Scale bar = 50 µm.

### Perturbation of miR-1 impacts neuronal development

Since gut contractions are in part mediated by serotonin (Yaguchi and Yaguchi, 2021), we examined the level of serotonin, number, and positioning of serotonergic neurons (Fig. 4A). Compared to control, miR-1 inhibitor-injected embryos had decreased levels of serotonin and fewer serotonergic neurons, while miR-1 mimic-injected embryos had a higher serotonin levels and more serotonergic neurons (Fig. 4Ai-ii). Additionally, we observed that serotonergic neurons in control-injected larvae were positioned linearly in the animal plate, whereas serotonergic neurons in miR-1 mimic-injected larvae are dispersed throughout the apical organ or within the ciliary band (Fig. 4Aiii). We also observed miR-1 perturbed embryos exhibited a significant decrease in mature and functional neurons compared to the control, as indicated by the immunolabeling of SynB-expressing neurons (Fig. 4B).

**Figure 4.**
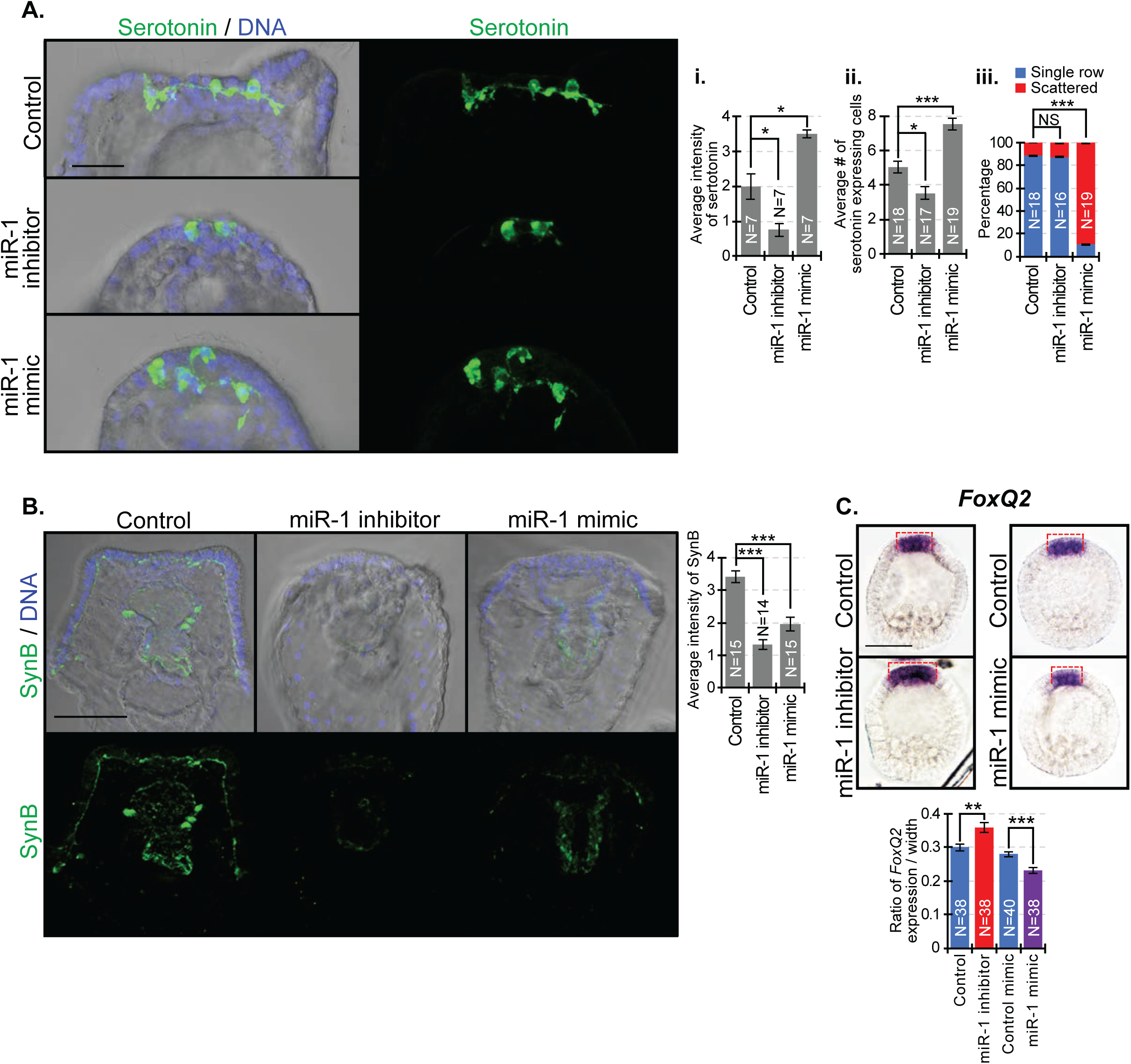
**Perturbation of miR-1 results in neuronal specification defects.** (A) Larvae were immunolabeled with Serotonin antibody (green) and counterstained with DAPI for DNA (blue). 4 biological replicates. (B) Larvae were immunolabeled with SynB antibody (green) and counterstained with DAPI (blue). (C) Blastulae were incubated with *FoxQ2* RNA probe. Compared to control, the expression domain of *FoxQ2* was significantly increased in miR-1 inhibitor-injected embryos, whereas miR-1 mimic-injections resulted in significantly decreased *FoxQ2* expression domain. 2 biological replicates. Student’s t-test. Scale bar = 50 µm.

Since FoxQ2 is required for serotonergic neural specification and must be downregulated for their differentiation (Yaguchi et al., 2012), we examined its expression. We observed that miR-1 inhibitor-injected embryos have expanded *FoxQ2* expression domain, while miR-1 mimic-injected embryos have decreased *FoxQ2* expression domain compared to control (Fig. 4C).

### miR-1 regulates pigment cell number and morphology

PCs are derived from the mesoderm and has an essential role in the innate immune defense system (Fig. 1A) (Tokuoka et al., 2002; Smith et al., 2006). In control embryos, PCs have long filopodial extensions (Fig. 5). In miR-1 inhibitor-injected larvae, PCs are clumped or rounded, whereas PCs in miR-1 mimic-injected larvae have multiple long filopodial extensions (Fig. 5Ai).

**Figure 5.**
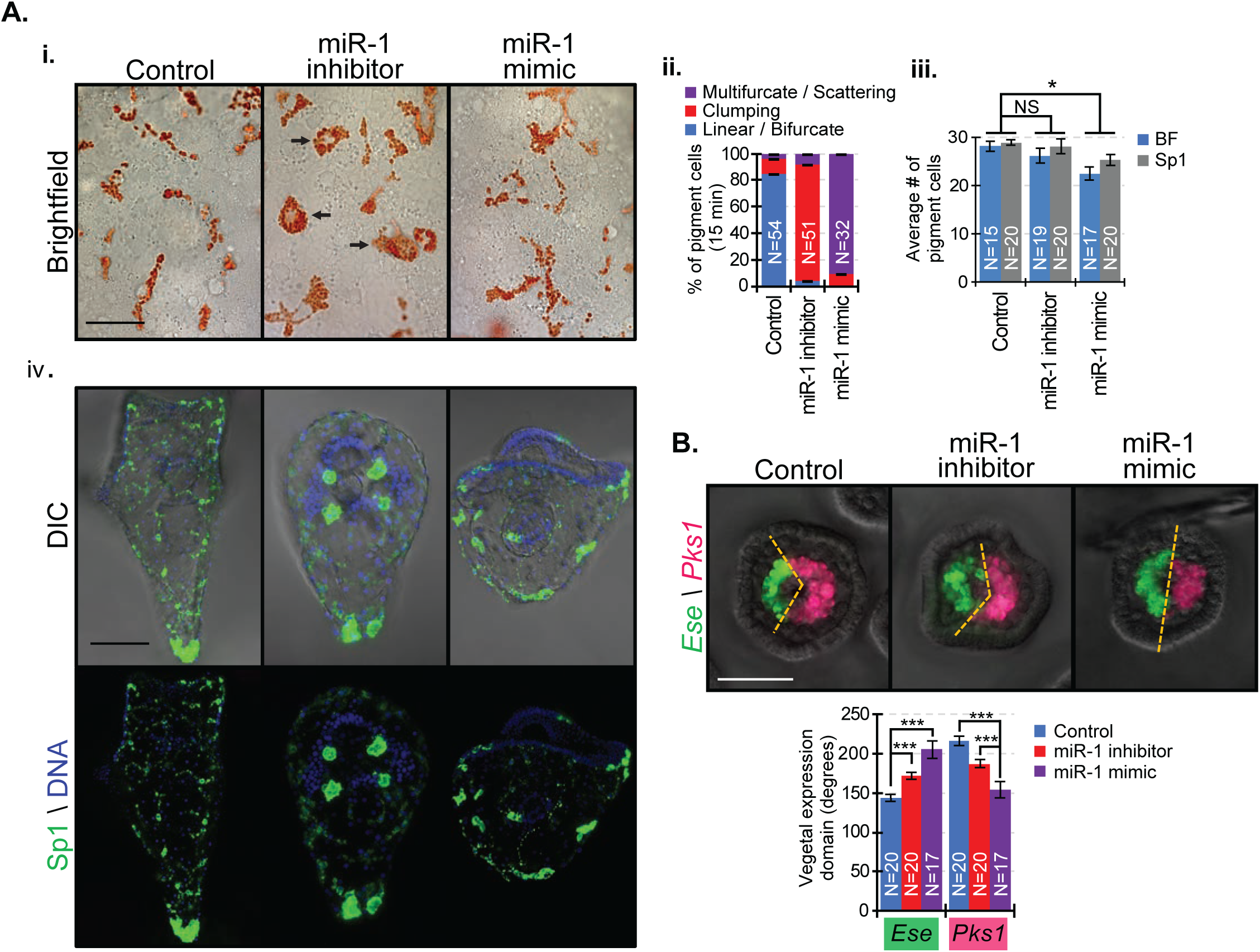
**Inhibition of miR-1 results in PC morphology changes**. (Ai) Snapshots of PCs in live larvae are depicted. Scale bar = 20 µm. (Aii) N=number of pigment cells observed in 7 embryos for each condition. Arrows indicate cells that lack filopodial extensions. 2 biological replicates. (Aiii) Compared to control, miR-1 mimic-injected embryos have fewer PCs, based on counting them in brightfield (BF) or with Sp1 immunolabeling. Student’s t-test. Scale bar = 50 µm. 2 biological replicates. (Aiv) Sp1 antibody was used to detect PCs. (B) F*IS*H was used to measure the vegetal spatial expression of BC TF *Ese* and PC marker *Pks1* during blastula stage, demarcated by the yellow dashed lines. Student’s t-test. 2 biological replicates.

In addition, while inhibition of miR-1 did not affect the number of PCs, miR-1 mimic-injections resulted in a significant decrease of PCs compared to control (Fig. 5Aii).

We further examined the behavior of PCs and observed that PCs of control larvae travel in a linear fashion with occasional bifurcated filopodial extensions (Fig. 5Ai, Movie 4). In miR-1 inhibitor-injected larvae, their rounded PCs move with membraneous blebs, whereas miR-1 mimic-injected larvae form multifurcated filopodial extensions that travel in multiple directions (Fig. 5Ai, Movies 5,6).

We examined the spatial expression of blastocoelar cell (BC) TF *Ese* (oral/ventral mesoderm) and pigment cell marker *Pks1* (aboral/dorsal mesoderm) in the mesenchyme blastula stage, when PCs and BCs become specified (Fig. 5B) (Materna et al., 2013). We observed that both inhibitor and miR-1 mimic-injections resulted in increased *Ese* and decreased *Pks1* expression domains, with miR-1 mimic-injected embryos having a more dramatic decrease in the expression domain of *Pks1* (Fig. 5).

### miR-1 perturbations result in less PMCs and a delay in PMC ingression

To assess how miR-1 perturbation impacts PMC specification and/or ingression, we used the *VegfR10* RNA probe to mark and follow PMCs during early gastrulation in a time-course experiment. While undergoing epithelial to mesenchymal transition (EMT), PMCs appear as ‘bottle-shaped’, whereas cells that have completed EMT have a ‘round cell shape’ (Lyons et al., 2012; Sampilo et al., 2021). Using the cell shape as a criterion for EMT, we found miR-1 perturbation resulted in a significant delay of PMC ingression during gastrulation (Fig. 6Ai,Bi). miR-1 perturbed embryos also had an average of three PMCs less than control embryo (Fig. 6Aii,Bii).

**Figure 6.**
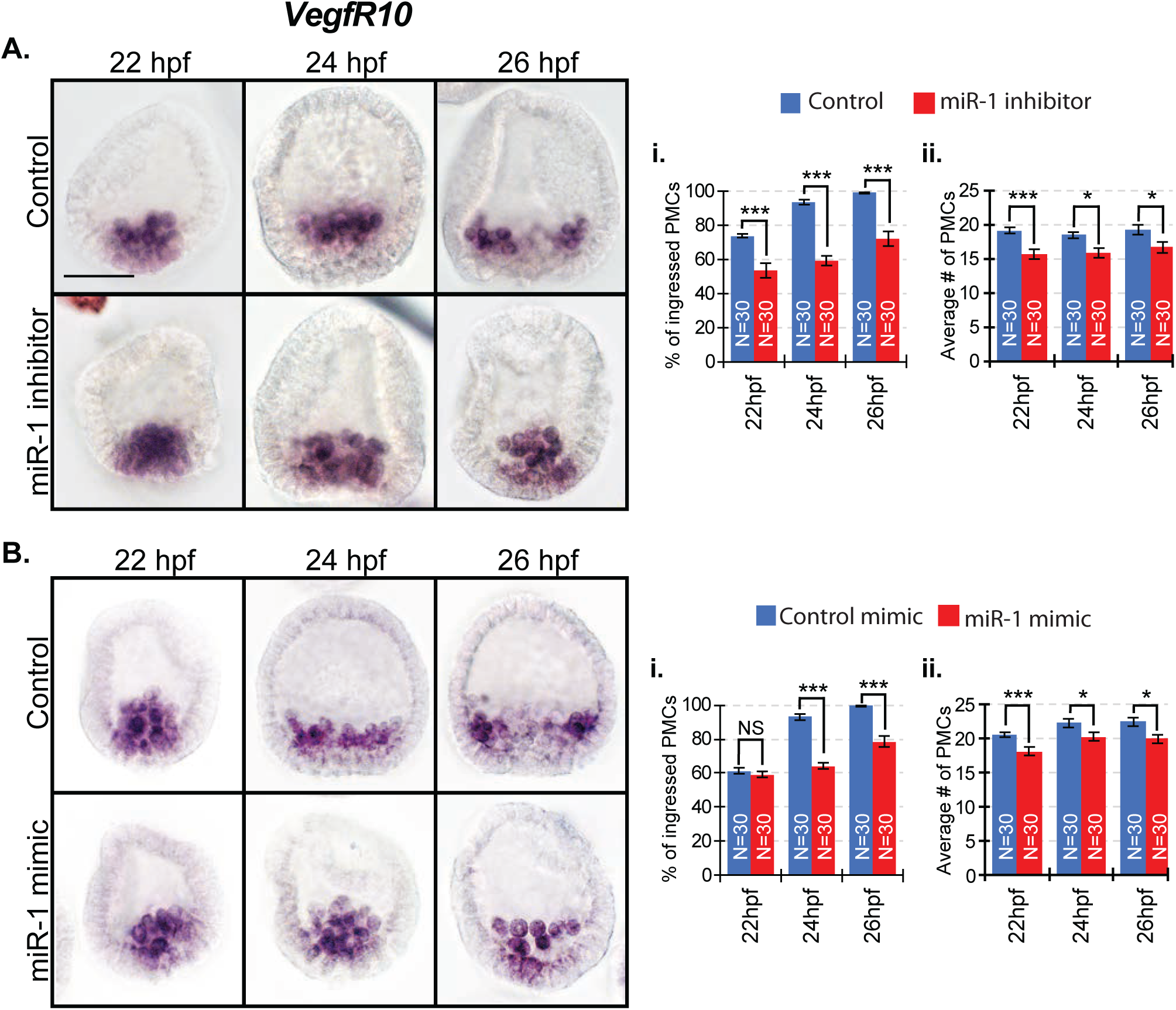
Perturbation of miR-1 induced a delay in PMC ingression. Control and miR-1 perturbed embryos were collected at various times during PMC ingression. (A) *VegfR10*-positive PMCs were counted through a series of Z-stack images to examine cell specification and EMT. Compared to control, miR-1 perturbed blastulae had less PMCs. Student’s t-test. (B) The percentage of “ingressed, round” PMCs from each embryo were calculated from the total of *VegfR10*-expressing PMCs (Sampilo et al., 2021). 3 biological replicates. Cochran-Mantel-Haenszel test. 3 biological replicates. Scale bar = 50 μm.

### Perturbation of miR-1 results in skeletal defects

During gastrula stage, we observed that miR-1 inhibitor and miR-1 mimic-injections resulted in decreased length of DVCs in a dose-dependent manner (Fig. 7A,B). miR-1 inhibitor-induced skeletal defects were rescued by co-injection of a miR-1 mimic, indicating that this defect is specifically induced by miR-1 inhibition (Fig. 7A). Interestingly, both miR-1 perturbations resulted in supernumerary tri-radiate rudiments and a similar ectopic skeletal branching off one of the tri-radiate arms (Fig. 7C).

**Figure 7.**
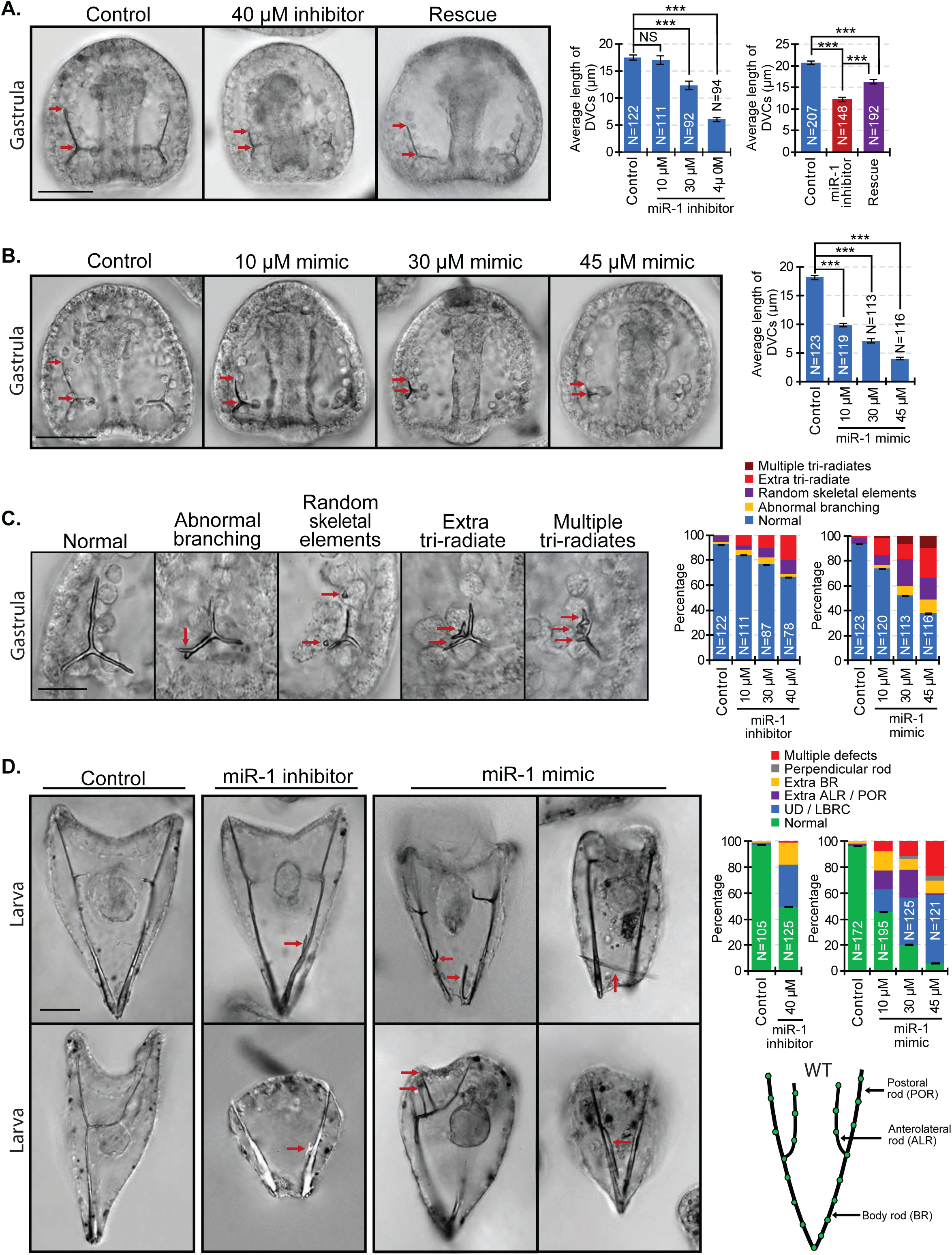
Perturbation of miR-1 results in shortening of the DVCs and abnormal skeletal branching. (A) miR-1 inhibitor was injected at various concentrations, resulting in decreased DVC length in a dose-dependent manner. Shortened DVCs can be rescued by co-injection of miR-1 inhibitor with the miR-1 mimic. Arrows indicate the length of DVCs. (B) miR-1 mimic-injected embryos had a dose-dependent decrease of DVCs. N is the total number of spicules examined. Student’s t-test. Scale bar = 50 μm. (C) Perturbation of miR-1 resulted in various skeletal defects (arrows), such as abnormal tri-radiate protrusions, scattered skeletal elements, and supernumerary tri-radiates. Scale bar = 20 µm. (D) miR-1 inhibitor-injected larvae exhibited ectopic BR branching, lack of BR convergence, and underdeveloped larvae. miR-1 mimic-injected embryos were underdeveloped and exhibited severe defects of supernumerary branching (arrows) off the PORs, ALRs, and/or BRs in a dose-dependent manner. 3 biological replicates. Scale = 50 μm.

miR-1 inhibitor-injected larvae were underdeveloped, lacked body rod (BR) convergence, and/or occasionally exhibited ectopic BR branching (Fig. 7D). miR-1 mimic-injected larvae had a dose-dependent severity of abnormal and supernumerary skeletal branching off the postoral rods (POR), anterolateral rods (ALR), and BRs (Fig. 7D). We also observed independent skeletal elements developed perpendicular to the larval BRs. Based on the dose-response data, 40 µM of miR-1 inhibitor and 30 µM of miR-1 mimic were used throughout this study. Overall, these results indicate that miR-1 plays a critical role in the initial formation and elongation of the skeletal spicules and that miR-1 mimic-injections induced a more severe skeletal defect than miR-1 inhibitor-injections.

### Perturbation of miR-1 results in PMC patterning defects

Since we observed skeletal branching defects (Fig. 7), we examined the patterning of PMCs, which are the only cells that make the skeleton (Ettensohn and McClay, 1986) (Fig. 8). In miR-1 inhibitor-injected gastrulae, we found that while the patterning of PMCs was not greatly affected, PMCs were clustered posteriorly and had less anterior migration compared to control (Fig. 8Ai-ii). This decreased migration in miR-1 inhibitor-injected embryos was rescued by co-injection of miR-1 mimic, indicating that this defect is due to inhibition of miR-1. In miR-1 mimic-injected gastrulae, the PMCs were dispersed and migrated to the animal pole (Fig. 8Bi-ii). miR-1-inhibitor-injected larvae had occasional PMCs positioned off a skeletal branch (Fig. 8Ci), whereas miR-1 mimic-injected larvae had many PMCs positioned off all branches (Fig. 8Cii).

**Figure 8.**
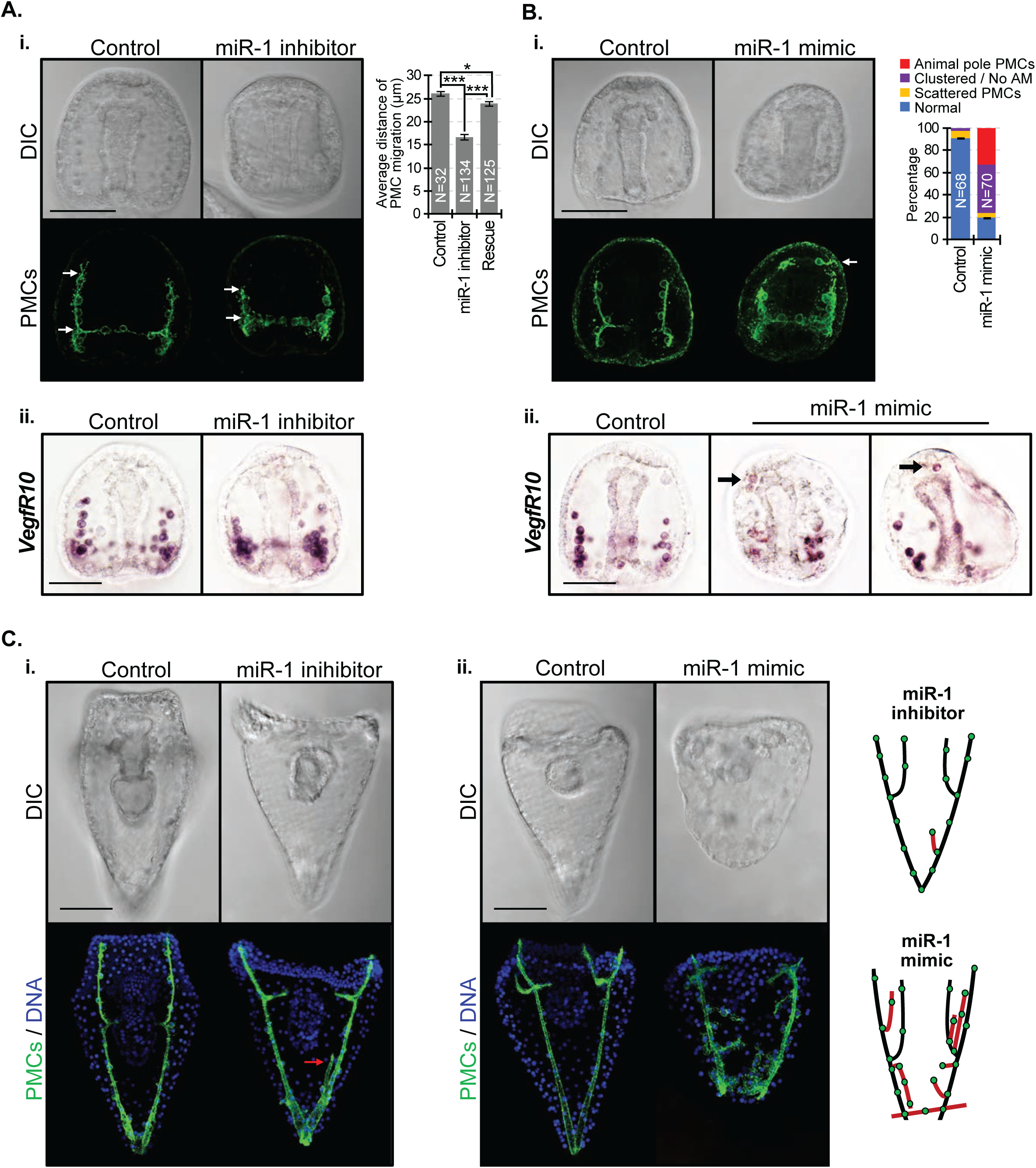
Perturbation of miR-1 results in ectopic PMC patterning. Gastrulae were immunolabeled with 1D5 PMC antibody (McClay et al., 1983) or hybridized with the *VegfR10* RNA probe. (Ai-ii) PMCs in miR-1 inhibitor-injected gastrulae exhibit less anterior migration (arrows) compared to the control that was able to be rescued by co-injection of miR-1 mimic. *VegfR10*-expressing cells also exhibit clustering and lack of anterior migration compared to control, similar to immunolabeling with the 1D5 antibody. Student’s t-test. 3 biological replicates. (Bi-ii) miR-1 mimic-injected gastrulae exhibited scattered or clustered PMCs, and PMCs that have migrated near animal pole (arrows). (C) Larvae were immunolabeled with 1D5 antibody and counterstained with DAPI (blue). (Ci) Compared to control, miR-1 inhibitor-injected larvae exhibited ectopic branching off the BR (arrow). (Cii) miR-1 mimic-injections resulted in severe PMC patterning defects, correlating to abnormal skeletal branching. Scale = 50 μm.

### miR-1 directly targets components of skeletogenic GRN and signaling pathways

To reveal the regulatory molecular mechanism of miR-1, we bioinformatically searched for potential miR-1 binding sites within transcripts that encode regulators of the PMC GRN and key components of signaling pathways. Using *R*luc reporter construct and site-directed mutagenesis, we identified miR-1 to directly repress *Ets1/2, Tbr, VegfR7, Notch, Nodal,* and *Wnt1.* miR-1 may have weak miRNA-mRNA binding affinity with *Dri* (Fig. 9A,B).

**Figure 9.**
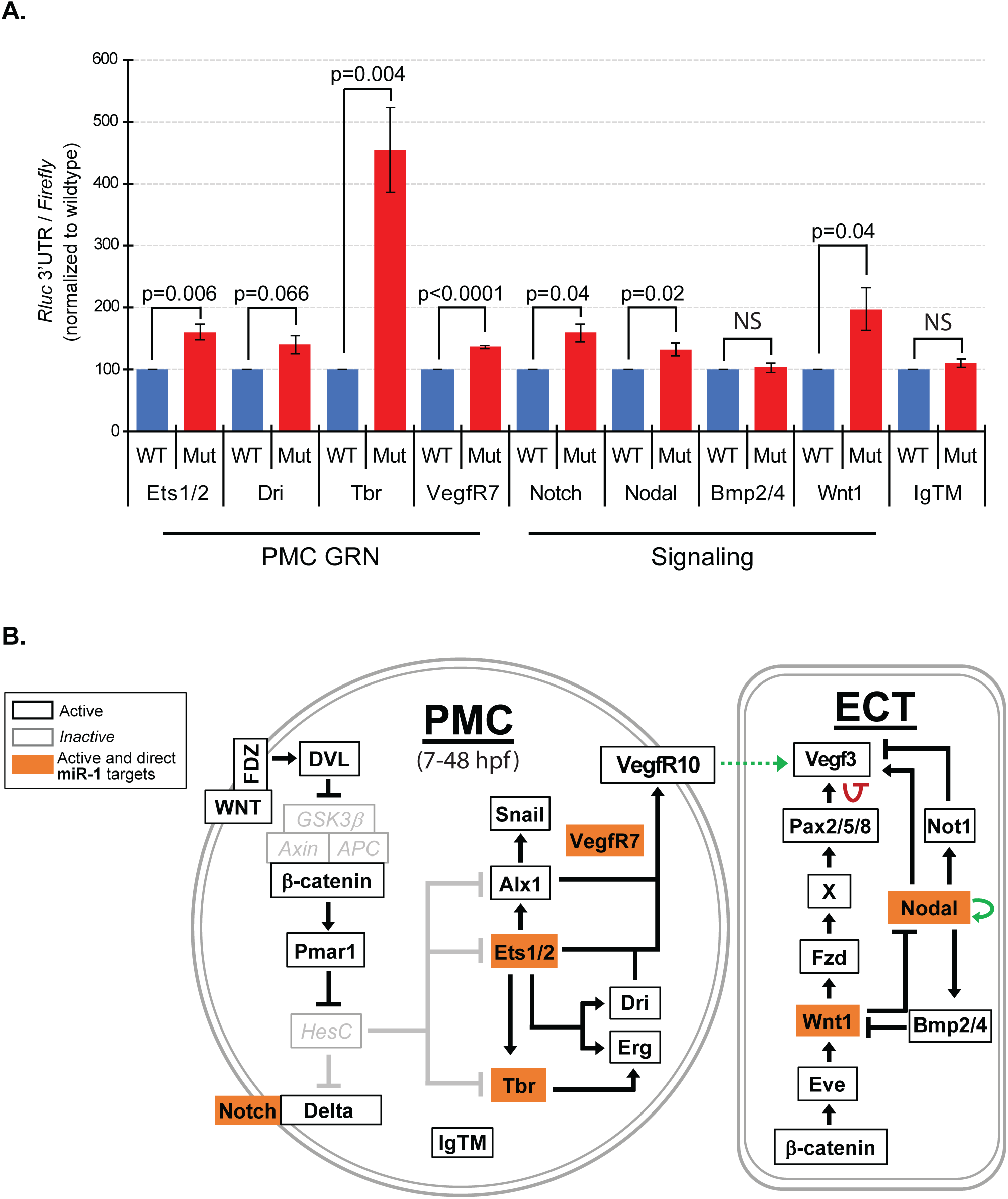
miR-1 targets multiple components skeletogenic GRN and signaling pathways. (A) miR-1 directly suppresses *Ets1/2, Tbr, VegfR7, Notch, Nodal* and *Wnt1*. Student’s t-test. (B) Schematic of simplified PMC GRN is shown. PMCs are specified by the Wnt/β-catenin signaling pathway which activates the transcriptional repressor *Pmar1,* leading to the activation of skeletogenic TFs *Ets1/2*, *Tbr*, *Alx1*, *Dri* and *Erg*, as well as other endodermally and mesodermally-derived TFs (not shown). In the ectoderm (ECT), Wnt/β-catenin signaling activates *Eve* which in turn activates *Wnt1*. Nodal in the ventral ectoderm activates BMP signaling to restrict *Wnt1* to the posterior-ventral side while Wnt1 prevents Nodal expression in the posterior region. Nodal activates *Not1,* and Not1 represses *Vegf3* in the ventral ectoderm, thus restricting *Vegf3* expression to the two lateral ectodermal domains. Validated miR-1 targets are highlighted in orange.

### miR-1 mimic-injections result in ectopic expression domains of several factors

To identify the molecular mechanism of how miR-1 regulates skeletogenesis, embryos were collected at gastrula stage and hybridized with *Vegf3, Wnt1, Nodal, Not1,* and *Bmp2/4* RNA probes (Fig. 10). In miR-1 inhibitor-injected gastrulae, the lateral and vegetal expression domains of *Vegf3* were significantly decreased compared to control (Fig. 10Ai). Concurrently, the vegetal expression domains of *Nodal* and *Not1* were expanded, without expression domain change of *Bmp2/4* (Fig. 10Bi). In miR-1 mimic-injected gastrulae, the lateral expression of *Vegf3* had no expression domain change; however, its vegetal expression domain was significantly expanded into both the VE and DE domains (Fig. 10Aii,Bii). In contrast to miR-1 inhibitor-injected embryos, the miR-1 mimic-injected embryos had *Nodal, Not1,* and *Bmp2/4* decreased in the vegetal expression domains, while *Wnt1* was increased compared to control. Results indicate that *Vegf3* expression domains in miR-1 perturbed embryos correlated to changes in expression domains of *Nodal*, *Bmp2/4*, and *Wnt1*.

**Figure 10.**
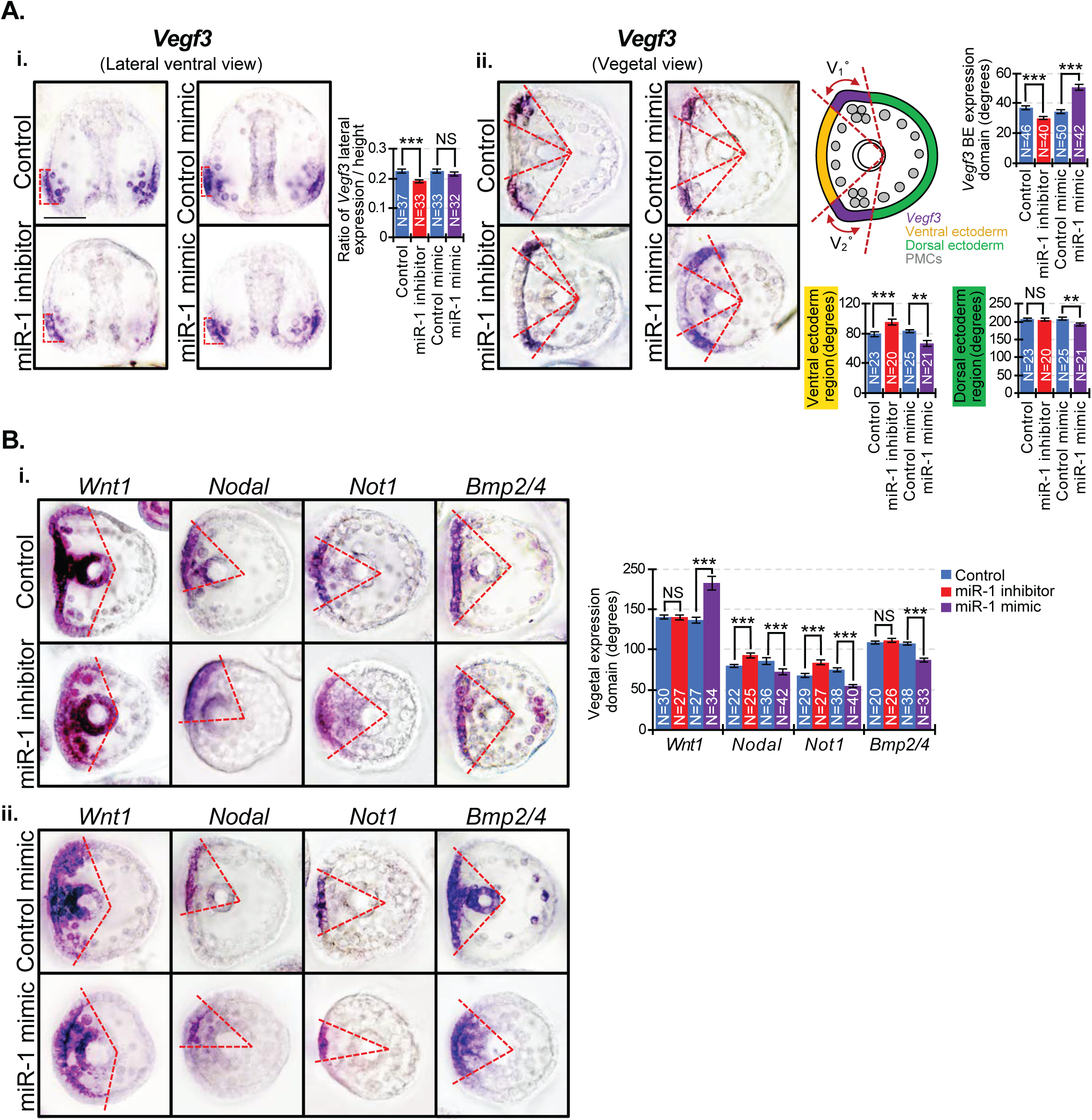
**Perturbation of miR-1 results in ectopic expression domains.** (A) Gastrulae were hybridized with *Vegf3, Wnt1, Nodal, Not1* and *Bmp2/4* RNA probes. (Ai) The lateral expression domain of *Vegf3* (red dashed lines) decreased in miR-1 inhibited gastrulae, while its ventral *Vegf3* expression domain decreased from the ventral ectoderm (VE; orange region in the schematic). (Aii) The lateral expression of *Vegf3* in miR-1 mimic-injections resulted in no change, while the vegetal expression domain of *Vegf3* in the border ectoderm (BE) was significantly expanded into the VE as well as into the dorsal ectoderm (DE; green region in the schematic). (Bi) In miR-1 inhibitor-injected gastrulae, *Nodal* and *Not1* vegetal expression domains expanded while the expression domains of *Bmp2/4* and *Wnt1* were not altered. (Bii) In miR-1 mimic-injections, *Nodal, Not1* and *Bmp2/4* vegetal expression domain decreased, while *Wnt1* and *Vegf3* expanded. N is the number of domains measured. Student’s t-test. 2-3 biological replicates. Scale = 50 μm.

### miR-1 regulates expression of key endodermal, ectodermal, and mesodermal transcripts

To identify the underlying molecular mechanism of how miR-1 impacts structures derived from the three germ layers, we tested expression levels of key TFs involved in endodermal, ectodermal, and mesodermal specification (Fig. 11A). Results indicate that miR-1 inhibition resulted in a 2-fold increase in *Krl* and *FoxA,* while *Onecut* decreased 2-fold. miR-1 mimic-injections resulted in almost a 2-fold increase in *Onecut,* and *Ese,* while *Tropo1, MHC,* and *Pks* had a >2-fold decrease.

**Figure 11.**
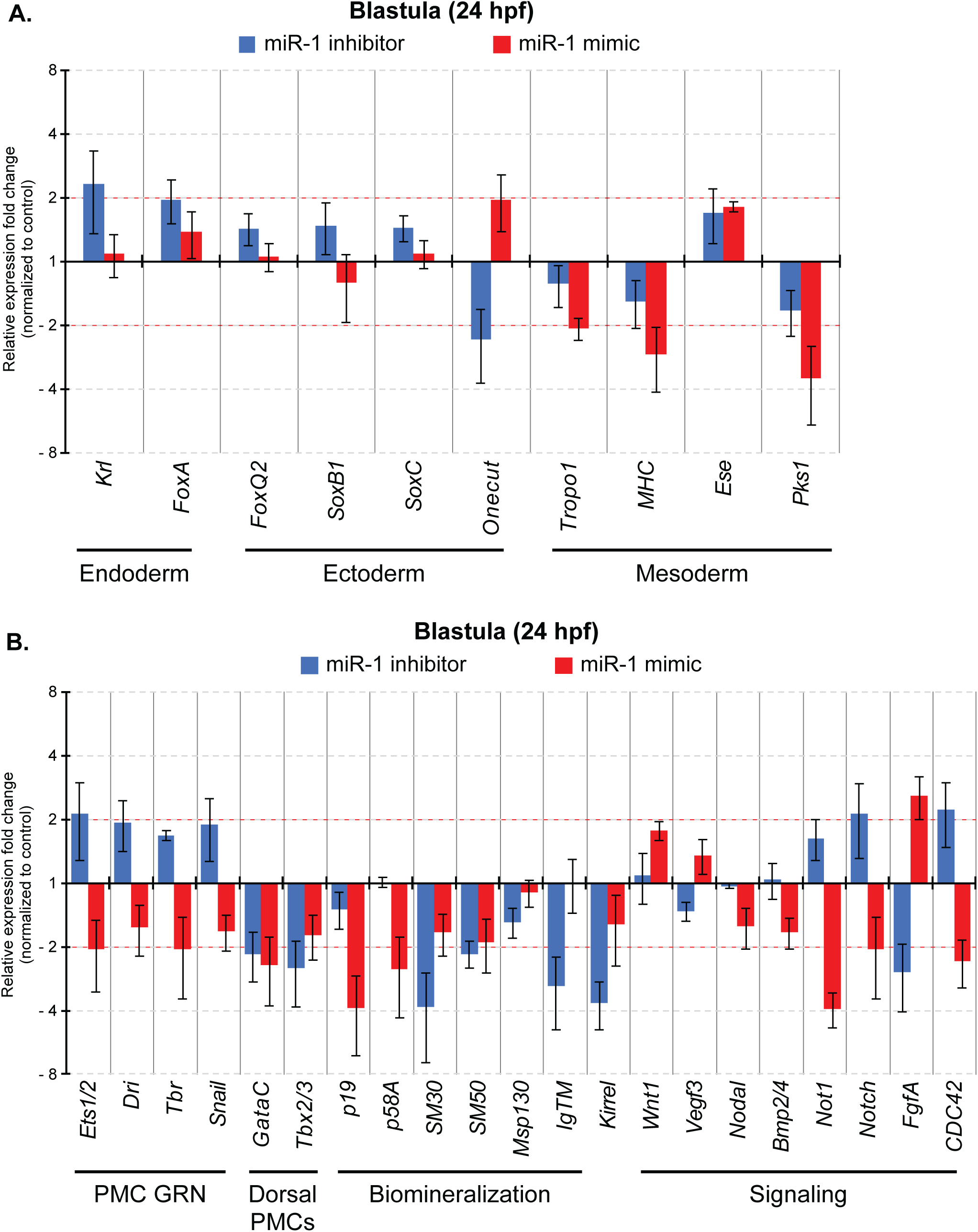
miR-1 regulates transcripts that encode key endodermal, ectodermal, and mesodermal TFs. (A) The expression levels of key TFs important in specification of the endoderm, ectoderm, and mesoderm were collected at the mesenchyme blastula stage and assessed with qPCR. (B) The expression levels of key transcripts encoding factors important for PMC specification and patterning were collected at the mesenchyme blastula stage and assessed with qPCR.

To identify the underlying molecular mechanism that led to PMC patterning and skeletal branching defects, we examined the relative expression levels of transcripts that encode PMC specification and patterning (*Ets1/2, Dri, Tbr, Snail, Nodal, Bmp2/4, Not1, Vegf3, Wnt1, FgfA* and *CDC42*) (Röttinger et al., 2004; Duloquin et al., 2007; Röttinger et al., 2008; Duboc et al., 2010; Adomako-Ankomah and Ettensohn, 2013; Sepúlveda-Ramírez et al., 2018), biomineralization enzymes (*P19, p58A, SM30, SM50*) (Cheers and Ettensohn, 2005; Livingston et al., 2006; Adomako-Ankomah and Ettensohn, 2011), PMC-specific cell surface protein *Msp130* (Leaf et al., 1987), and PMC adhesion protein *KirrelL* (Ettensohn and Dey, 2017), and markers of dorsal PMCs (*Tbx2/3* and *GataC*) (Duboc et al., 2010) (Fig. 11B). Results indicate that inhibition of miR-1 results in a 2-fold increase in transcript levels of *Ets1/2, Notch,* and *CDC42*, while miR-1 mimic-injections resulted in decreased transcript levels of these same genes. Inhibition of miR-1 resulted in an almost 2-fold increase in *Dri, Tbr,* and *Snail*. Both miR-1 inhibitor and miR-1 mimic-injections resulted in ∼2-fold decrease in *GataC, Tbx2/3, SM30,* and *SM50*.

## Discussion

We identified miR-1 to have a diverse role in embryogenesis, in addition to its conserved function in muscle-specific tissues. In the sea urchin, miR-1 is one of the most abundantly expressed miRNAs, and was able to rescue severe embryonic lethality induced by knockdown of Drosha and/or Dicer when co-injected with the three other highly sequenced miRNAs in the sea urchin (miR-31, miR-71, and miR-2012) (Song et al., 2012), suggesting that miR-1 is required for proper early development. We identified miR-1 to not only regulate mesodermally-derived structures, which include muscles, PCs, and skeletogenesis, but also gut morphology and function, and specification and positioning of serotonergic neurons.

We observed that the gut morphology and contractions in miR-1 perturbed larvae were altered compared to control (Fig. 2Bi-iii,2C, Movies 1-3). Both miR-1 inhibitor and mimic-injected embryos have expanded expression domains of *Krl* and *FoxA* (Fig. 2B-D). Previously, *Krl* knockdown resulted in missing foregut and delayed gastrulation (Howard et al., 2001; Yamazaki et al., 2008), while *FoxA* knockdown reduces gut size (Oliveri et al., 2006). Consistent with our *in situ* data, miR-1 inhibition resulted in a 2-fold increase of *Krl* and *FoxA* (Fig. 11A). Since *Krl* and *FoxA* do not contain canonical miR-1 binding sites, miR-1 may indirectly modulate them to impact gut development. In addition, we found that miR-1 directly suppresses *Notch* and *Wnt1* (Fig. 9). Consistent with prior work, *Notch2* is a known miR-1 target (Liu et al., 2018; Hua et al., 2019). *Notch* knockdown embryos resulted in decreased *Krl* levels (Yamazaki et al., 2008), and *Wnt1* knockdown resulted in decreased *FoxA* levels (Cui et al., 2014). Thus, miR-1 inhibitor-injected embryos may have increased Notch and Wnt1, which may result in increased *Krl* and *FoxA*. While *Krl* and *FoxA* expression domain expanded in miR-1 mimic-injected embryos, their levels did not increase (Figs. 2D, 11A). How miR-1 mimic-injection resulted in expanded *Krl* and *FoxA* is unclear. miR-1 may also regulate transcripts that encode other undefined factors important for endodermal differentiation, gut morphology, and function.

The defective gut contractions in miR-1 perturbed larvae may be due to problems with the muscle fibers and/or neural coordination. We observed that miR-1 perturbed larvae have decreased number of F-actin muscle fiber rings (Fig. 3). Previous work has shown Delta/Notch signaling to play critical roles in vertebrate myogenesis (Kopan et al., 1994; Conboy and Rando, 2002; Schuster-Gossler et al., 2007), and knockdown of either Delta or Notch in the sea urchin embryos resulted in decreased muscle fibers (Sweet et al., 2002; Range et al., 2008). We showed that miR-1 directly suppresses *Notch*, indicating that the defective muscle fibers in miR-1 perturbed larvae may be in part explained by miR-1’s direct suppression of *Notch* (Fig. 9A). Additionally, we bioinformatically identified miR-1 to have predicted truncated and/or mismatched miR-1 binding sites within *MHC* and *Tropo1*, which are conserved muscle differentiation genes (Mu et al., 2007; Andrikou et al., 2013; Annunziata et al., 2014; Jin et al., 2016). In the sea urchin, both are expressed in circumesophageal muscle fibers, and *MHC* is also expressed in all the sphincters (Andrikou et al., 2013; Annunziata et al., 2014). In vertebrates, various MHC isoforms are expressed in the upper esophageal sphincter and have distinct contractile properties (Mu et al., 2007). The exact function of Tropo1 in the sea urchin is unknown; however, in vertebrates, Tropo1 is critical in cardiac development and contractile function of skeletal muscles (Jin et al., 2016; England et al., 2017). While the levels of *MHC* and *Tropo1* transcripts in miR-1 inhibitor-injected embryos did not change compared to the control, their levels were decreased 2-fold in miR-1 mimic-injected embryos (Fig. 11A). Interestingly, miR-1 is enriched in the larval gut (Fig. 2B), suggesting that it may post-transcriptionally regulate *MHC* and *Tropo1* that may impact the organization of the F-actin of the muscle fibers and/or gut contractions. Additionally, since serotonin can regulate the pyloric opening of the larval gut (Yaguchi and Yaguchi, 2021), aberrant serotonin expression in miR-1 perturbed embryos may also contribute to decreased gut contractions (Figs. 2C, 4A).

We found that in comparison to control larvae, miR-1 inhibitor-injected larvae had lower serotonin levels and fewer serotonergic neurons, while miR-1 mimic-injected larvae had increased serotonin levels and a higher number of serotonergic neurons (Fig. 4A). We propose that miR-1 regulates the specification of serotonergic neurons, potentially through *FoxQ2*. FoxQ2 is required early for the specification of serotonergic neurons and must be removed from the apical plate by late gastrula stage to allow differentiation of serotonergic neurons (Yaguchi et al., 2008; Yaguchi et al., 2012). In miR-1 inhibitor-injected blastulae, the expression domain of *FoxQ2* is expanded (Fig. 4C), consistent with prior findings where expanded *FoxQ2* in LRP6-morphants correlated with a complete loss of serotonergic neurons (Yaguchi et al., 2016). In miR-1 mimic-injected blastulae, *FoxQ2* expression domain became more restricted which may result in increased serotonergic neurons (Figs. 4A,C). It appears that while the spatial expression of *FoxQ2* was altered in miR-1 perturbed embryos, *FoxQ2* levels did not change (Fig. 11A). Since *FoxQ2* does not contain potential miR-1 binding sites, we speculate that miR-1 may indirectly regulate expression of *FoxQ2* in early development.

miR-1’s regulation of *FoxQ2* may be through its direct suppression of *Nodal*. (Fig. 9). Nodal has been implicated in promoting neuronal specification in humans and in sea urchins (Smith et al., 2008; Patani et al., 2009; Yaguchi et al., 2016; McClay et al., 2018). In *Nodal* morphants, *FoxQ2* expression domain in blastulae became more restricted (Yaguchi et al., 2016), and this restriction of *FoxQ2* expression domain is similar to what was observed in miR-1 mimic-injected blastulae (Fig. 3C). Thus, miR-1 may suppress *Nodal* to indirectly restrict *FoxQ2*. Interestingly, serotonergic neurons of *Nodal* morphants were dispersed throughout the apical domain (Yaguchi et al., 2016), similar to what we observed in miR-1 mimic-injected larvae (Fig. 3A). Nodal/Bmp signaling along the DV axis is critical for regulating the specification of the neuroectoderm and patterning within the apical domain (Duboc et al., 2004; Yaguchi et al., 2016). In the sea urchin, Nodal signaling suppresses serotonergic neuronal differentiation from the oral/ventral side of the animal plate (Yaguchi et al., 2007; Yaguchi et al., 2016). Thus, potentially, aberrant Nodal expression induced by miR-1 overexpression may lead to mispositioning of these serotonergic neurons.

Additionally, we found that miR-1 perturbation resulted in a significant decrease in SynB-positive neurons compared to the control (Fig. 4B). SynB mediates neurotransmitter release of synaptic vesicles in mature and functional neurons (Burke et al., 2006; Yang et al., 2015). Since neuronal TFs *SoxB1* and *SoxC,* which regulate neuronal specification (Angerer et al., 2005; Garner et al., 2016; Niwa et al., 2016), were not significantly altered under miR-1 perturbation (Fig. 11A), this indicates that miR-1 may regulate neuronal development by suppressing *Nodal*, and indirectly *FoxQ2*, as well as other TFs during neuronal differentiation.

We then examined mesodermally-derived structures in miR-1 perturbed embryos. In the sea urchin, PCs migrate throughout the larval ectoderm to carry out immune defense (Takata and Kominami, 2003; Smith et al., 2006). miR-1 mimic-injected larvae have fewer PCs, correlating to decreased expression of *Pks1* (Figs. 5, 11A). During development, PCs and BCs share the same progenitor cells and make a binary fate decision during the blastula stage (Sweet et al., 1999). Their specification is regulated by both Nodal and Notch signaling pathways, where Nodal initiates specification by activating TFs that specify BCs and inhibiting TFs that specify PCs; later Notch signaling promotes differentiation of both BCs and PCs (Duboc et al., 2010). *Nodal* overexpression resulted in a lack of *Pks1-*expressing PCs and expanded expression of BC markers *GataC* and *Ese* (Duboc et al., 2010). Knockdown of *Notch* resulted in fewer PCs and BCs (Sweet et al., 1999; Peterson and McClay, 2005). Taken together, miR-1’s regulation of both *Nodal* and *Notch* is likely to contribute to BC and PC specification. PCs in miR-1 perturbed larvae exhibited different morphology and behavior when compared to control (Fig. 5, Movies 4-6). How miR-1 regulates the morphology and movement of PCs remains unclear; however, miR-1 perturbation affects the expression of CDC42, which may be important for mediating general cell movement (Fig. 11) (Yamao et al., 2015; Sepúlveda-Ramírez et al., 2018).

Another mesodermally-derived tissue that is regulated by miR-1 is the larval skeleton. To understand how miR-1 may be regulating PMC development and skeletogenesis, we identified miR-1 to directly suppress *Ets1/2, Tbr,* and *VegfR7* of the PMC GRN, and *Notch, Nodal, Bmp2/4,* and *Wnt1* of signaling pathways (Fig. 9). Ets1/2 and Tbr are critical for PMC specification and function. Ets1/2 is an important modulator of MAPK pathway that is required for the specification of PMCs and their ability to undergo EMT (Kurokawa et al., 1999; Röttinger et al., 2004). Additionally, Tbr plays an essential role in specification of the skeletogenic mesoderm and formation of the larval skeleton where *Tbr* knockdown resulted in a complete loss of skeleton (Oliveri et al., 2002). The percentage of PMCs undergoing EMT is significantly and consistently less than the control (Fig. 6Ai,6Bi). Additionally, we observed both miR-1 perturbed embryos have a decrease of 3-4 PMCs compared to control which would likely not have detrimental effects (Fig. 6Aii,6Bii). Thus, miR-1’s regulation of *Ets1/2* and *Tbr* may potentially contribute to the decreased PMCs and EMT defect in miR-1 perturbed embryos. Moreover, the delay in PMC ingression could be due to miR-1’s potential regulation of *G-cadherin* and *Snail,* both of which have predicted miR-1 binding sites. *Snail* represses the transcription of cadherin in order for PMCs to detach from basal lamina and ingress into the blastocoel (Wu and McClay, 2007). The delay in PMC ingression could potentially affect PMC pattering by disrupting the time and distance-sensitive interaction between *VegfR10*-expressing PMCs and the *Vegf3* ligand expressed in the ectoderm, as *Vegf3* expression becomes restricted to the Veg1 ectoderm by 30 hpf (Li et al., 2014). Additionally, the function of another Vegf receptor, VegfR7 is not well understood; however we have previously found that blocking miR-31’s direct suppression of *VegfR7* resulted in mislocalized PMCs, indicating VegfR7’s potential role in PMC patterning (Stepicheva and Song, 2015). Thus, miR-1’s direct suppression of *Ets1/2, Tbr,* and *VegfR7* may affect not only PMC ingression but also affect PMC patterning.

A possible mechanism for miR-1 perturbation-induced ectopic branching can be explained by our observation that the PMCs of miR-1 perturbed larvae migrate off the main skeletal elements and continue to form syncytial cables (filopodia) (Fig. 8C). Since PMCs are solely responsible for the formation of the larval skeleton (Ettensohn and McClay, 1986), these PMCs in miR-1 perturbed larvae appear to be mispatterned to areas where we observed ectopic skeletal elements (Figs. 7D, 8C). Additionally, ectopic branching can also be explained by miR-1’s direct suppression of *Nodal* (Fig. 9B). Of note is that miR-1 mimic-injected embryos have the characteristic radialized morphology in the vegetal view, similar to *Nodal* morphant-induced morphology (Figs. 10Aii,10Bii) (Duboc et al., 2010). Also, although the level of *Nodal* transcripts was not greatly altered in miR-1 mimic-injected embryos, we observed a >3-fold decrease of Nodal target, *Not1* (Li et al., 2012). This result indicates that miR-1 mimic-injected embryos have decreased Nodal protein, resulting in decreased *Not1* transcripts (Fig. 11B). miR-1’s direct suppression of *Nodal* could potentially explain the skeletal defects in both contexts of loss-of-function and gain-of-function of miR-1 (Figs. 7,9), since knockdown or overexpression of Nodal results in ectopic skeletal branching (Duboc et al., 2004). Additionally, Nodal and Bmp2/4 restrict *Fgfa* expression to the PMC VLC (Guss and Ettensohn, 1997; Röttinger et al., 2008). Our results indicate that miR-1 inhibition led to increased Nodal, which may lead to a >2-fold decrease of *Fgfa*; while miR-1 mimic-injections may lead to decreased Nodal which may lead to a >2-fold increase of *Fgfa* (Fig. 11B). During the gastrula stage, *Fgfa* is expressed in ectoderm near the regions of the tri-radiate spicule rudiment branches which form the BRs and the PORs (Horstadius, 1973; Röttinger et al., 2008). Inhibition of *Fgfa* prevents skeletogenesis, whereas overexpression of *Fgfa* resulted in supernumerary tri-radiates leading to ectopic skeletal branching. Increased *Fgfa* in miR-1 mimic-injected embryos could in part explain the multiple spicules in gastrulae and ectopic BR and POR branches observed in larvae (Fig. 7C,D). Interestingly, *Bmp2/4* knockdown also result in larvae with multiple skeletal rods and ectopic branching (Duboc et al., 2004). Since results indicate that miR-1 does not directly regulate *Bmp2/4*, any changes in *Bmp2/4* spatial expression and/or transcript levels are likely due to miR-1’s suppression of *Nodal* which activates *Bmp2/4* (Duboc et al., 2004). Thus, we propose that mispatterned PMCs and aberrant skeletal branching in miR-1 mimic-injected embryos may attribute to miR-1’s repression of *Nodal* to increase *Fgfa* (Duboc et al., 2010) and miR-1’s indirect regulation of *Bmp2/4*.

To further elucidate the molecular mechanism of how miR-1 regulates PMC patterning, we examined the spatial expression domains and transcript levels of *Vegf3, Wnt1, Nodal, Not1,* and *Bmp2/4* (Figs. 10,11). Both Nodal and Bmp2/4 signaling pattern the PMCs by restricting *Vegf3* expression to the two lateral ectodermal domains and posterior ventral corners where the PMC clusters reside to form the skeletal rudiments within the border ectoderm-dorsal ventral margin (BE-DVM) intersection (Fig. 12A) (Duboc et al., 2010; McIntyre et al., 2013; Layous, 2020). In the ventral ectoderm, Nodal activates the expression of itself, *Not1, Bmp2/4* and *Vegf3* at 8-9 hpf in the ventral ectoderm (Lapraz et al., 2009; Li et al., 2012). Subsequently, Not1 restricts *Vegf3* expression from the ventral domain by 27 hpf (Li et al., 2012), while Bmp2/4 protein translocates to the dorsal side by late blastula to eliminate *Wnt1* from the dorsal posterior cells to induce dorsal fates (Lapraz et al., 2009; Wei et al., 2012). Wnt1 then restricts *Nodal* from the posterior ventral region (Wei et al., 2012), and indirectly activates *Vegf3* (Sampilo et al., 2021). In miR-1 inhibitor-injected gastrulae, both the lateral and vegetal expression of *Vegf3* was significantly decreased (Fig. 10A). In contrast, miR-1 mimic-injected gastrulae had no change in lateral *Vegf3* expression (Fig. 10Ai), but vegetal *Vegf3* expression had significantly expanded into both the VE and DE domains (Fig. 10Aii). The expanded *Vegf3* expression domain may contribute to ectopic supernumerary skeletal branching (Duloquin et al., 2007; Morgulis et al., 2019). Previous studies in vertebrates have shown miR-1 to directly target *VegfA* (Stahlhut et al., 2012; Zhu et al., 2018). In the sea urchin, a partial *Vegf3* transcript contains one predicted miR-1 binding site; however, the sequence data on *SpVegf3* 3’UTR is incomplete and difficult to clone. Thus, although the change in *Vegf3* expression domain may be due to miR-1’s direct suppression, our results indicate that the expression of *Vegf3* in the ventral-dorsal axis may be a result of miR-1’s direct regulation of *Nodal*. Additionally, a previous study has shown that a loss of *Bmp2/4* resulted in the position of the PMC ring to shift towards the animal pole and a radialized the embryo (Angerer et al., 2000; Layous, 2020). Interestingly, in miR-1 mimic-injected gastrulae, we observed mispatterned PMCs that migrated near the animal pole (Fig. 8B), decreased *Bmp2/4* expression domain, and radialized embryos (Fig. 10Bii), indicating that these defects are likely due to miR-1’s suppression of *Nodal*. Animal pole PMCs that ectopically migrated from the DVCs may become PMCs that give rise to ectopic branches off the ALRs (Sun and Ettensohn, 2014).

**Figure 12.**
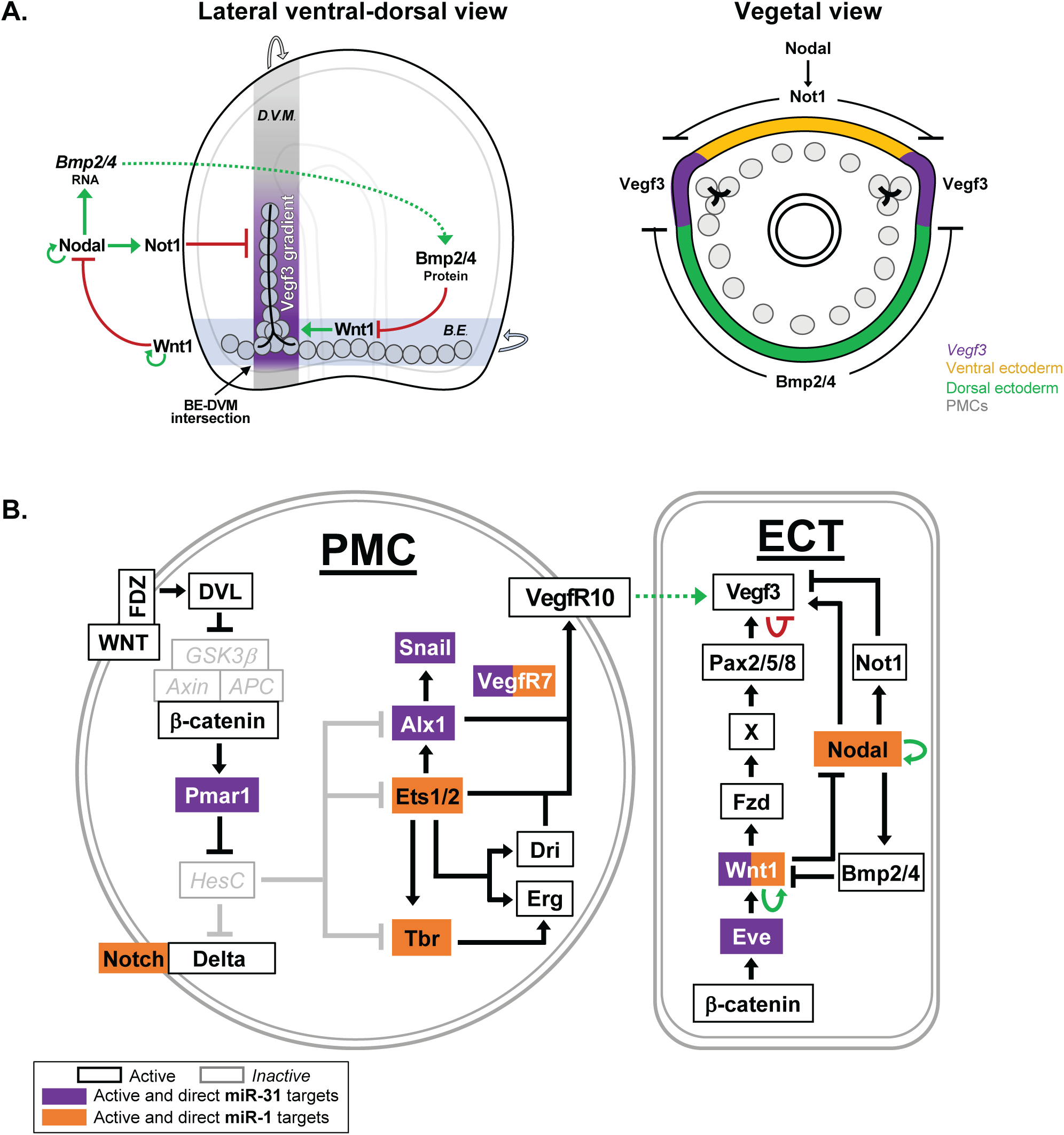
miR-1 and miR-31 regulate the expression of PMC GRN. (A) Nodal signaling activates its own transcription within the ventral ectoderm, which in turn activates *Bmp2/4* and *Not1.* Bmp2/4 ligand is translocated to the dorsal side of the embryo to activate signaling in the dorsal PMCs. Bmp2/4 indirectly restricts *Vegf3* ventrally by repressing Wnt1 that is upstream of *Vegf3* expression in the BE-DVM (border ectoderm dorsal-ventral margin). Nodal activation of *Not1* restricts *Vegf3* expression to the two lateral ectodermal domains. BE-DVM intersection is where the PMC clusters reside to form the skeletal rudiments (Duboc et al., 2010; McIntyre et al., 2013; Layous, 2020). (B) miR-1 directly suppresses transcripts (orange) such as *Ets1/2, Tbr* and *Notch* which are downstream of miR-31 target*, Pmar1* (purple). Both miR-1 and miR-31 directly suppress *VegfR7* and *Wnt1* (Stepicheva and Song, 2015; Sampilo et al., 2021). miR-1 and miR-31 are critical for proper PMC development.

The expanded expression domain of *Wnt1* in miR-1 mimic-injected embryos may restrict the expression domain of *Nodal*, leading to expansion of *Vegf3* expression domain. Although *Wnt1* is directly suppressed by miR-1, Wnt1 can activate its own transcription and is part of highly cross-regulated positive feedback circuitry (Cui et al., 2014; Sampilo et al., 2021). The expanded expression domain of *Wnt1* in miR-1 mimic-injected gastrulae is likely due to decreased *Bmp2/4* expression domain via miR-1’s direct suppression of *Nodal* (Figs. 9, 10Bii, 11B). The changes we observed in the spatial expressions of *Vegf3, Nodal, Not1, Bmp2/4,* and *Wnt1* in miR-1 mimic-injected embryos may all contribute to PMC mispatterning (Figs. 8B, 8Cii, 10Aii, 10Bii). Overall, miR-1 mimic-injected embryos have reminiscent larval branching defects as *Nodal* and *Bmp2/4* knockdown embryos (Duboc et al., 2004), and *Vegf3* and *Fgfa* overexpressing embryos (Fig. 7D) (Duloquin et al., 2007; Röttinger et al., 2008).

Previously, we identified that miR-31 in the sea urchin regulates skeletogenesis by directly suppressing *Eve* and *Wnt1* (Sampilo et al., 2021). Depletion of miR-31 resulted in expanded vegetal spatial expression of *Vegf3* (Stepicheva and Song, 2015; Sampilo et al., 2021), similar to miR-1 mimic-injections. miR-1 directly suppresses genes *Ets1/2, Tbr, VegfR7* and *Wnt1,* of which *Ets1/2* and *Tbr* are downstream of miR-31 targets *Pmar1* and *Eve* (Stepicheva and Song, 2015; Sampilo et al., 2021). Thus, both miR-31 and miR-1 target critical components within the PMC GRN and co-regulate skeletogenesis (Fig. 12B).

Overall, we identified miR-1 to regulate not only mesodermally-derived structures, but also mediate the development of the endodermal gut and ectodermally-derived nervous system. This study identifies novel functions of miR-1, by identifying its direct targets and revealing miR-1 as a multifunctional miRNA that regulates structures derived from all three germ layers in a developing embryo.

## Abbreviations

ALP: alkaline phosphatase
ALR: anterolateral rod
BC: blastocoelar cells
BR: body rod
BE-DVM: border ectoderm-dorsal ventral margin
cWnt: canonical Wnt
DE: dorsal ectoderm
Dpf: days post fertilization
DVC: dorsoventral connecting rod
EMT: epithelial to mesenchymal transition
GRN: gene regulatory network
PC: pigment cells
Hpf: hours post fertilization
PC: pigment cell
PMCs: primary mesenchyme cells
POR: postoral rod
TF: transcription factor
VE: ventral ectoderm
Vegf: vascular endothelial growth factor

## Acknowledgements

The authors would like to thank David McClay (Duke University) for his kind gift of 1D5 antibody and Gary Wessel (Brown University) for the SynB antibody. This work is funded by University of Delaware Graduate Fellowship to NFS. NSF CAREER (IOS 1553338) and MCB 2103453 to JLS and NIH NIGMS P20GM103446.

## Supplementary Figures

**Fig. S1 miR-1 is evolutionarily conserved.** Alignment of mature miR-1 sequences in several metazoan species is shown. Red nucleotides indicate the highly conserved miR-1 seed sequence. miR-1 in protostomes is shown in blue and deuterostomes in purple. Yellow indicates variable nucleotides. miR-1 sequences were obtained from miRBase (Kozomara and Griffiths-Jones, 2014) and aligned with Clustal Omega Multiple Sequence Alignment program (Sievers et al., 2011).

**Movie 1. Gut contractions in control embryos.** Live embryos were collected 5 dpf and recorded. Average of contractions was analyzed within 4 min. Scale bar = 20 µm.

**Movie 2. Gut contractions in miR-1 inhibitor-injected embryos.** Live embryos were collected 5 dpf and recorded. Average of contractions was analyzed within 4 min. Scale bar = 20 µm.

**Movie 3. Gut contractions in miR-1 mimic-injected embryos.** Live embryos were collected 5 dpf and recorded. Average of contractions was analyzed within 4 min. Scale bar = 20 µm.

**Movie 4. Pigment granules of control-injected larvae travel linearly.** Live embryos were collected 5 dpf and recorded. Pigment granules within the PCs of control larvae travel in a linear fashion that occasionally bifurcate into two filopodial extensions. Scale bar = 20 µm.

**Movie 5. Pigment cells in miR-1 inhibited larvae move with blebbing.** Live embryos were collected 5 dpf and recorded. In miR-1 inhibitor-injected larvae, pigment granules within immunocytes migrate in clumped or rounded manner. Scale bar = 20 µm.

**Movie 6. Pigment granules of miR-1 mimic-injected larvae exhibit scattered and multifurcated extensions.** Live embryos were collected 5 dpf and recorded. In miR-1 mimic-injected larvae, PCs exhibit scattered pigment granules that form multifurcated filopodial extensions that travel in multiple directions. Scale bar = 20 µm.

**Table S1.**
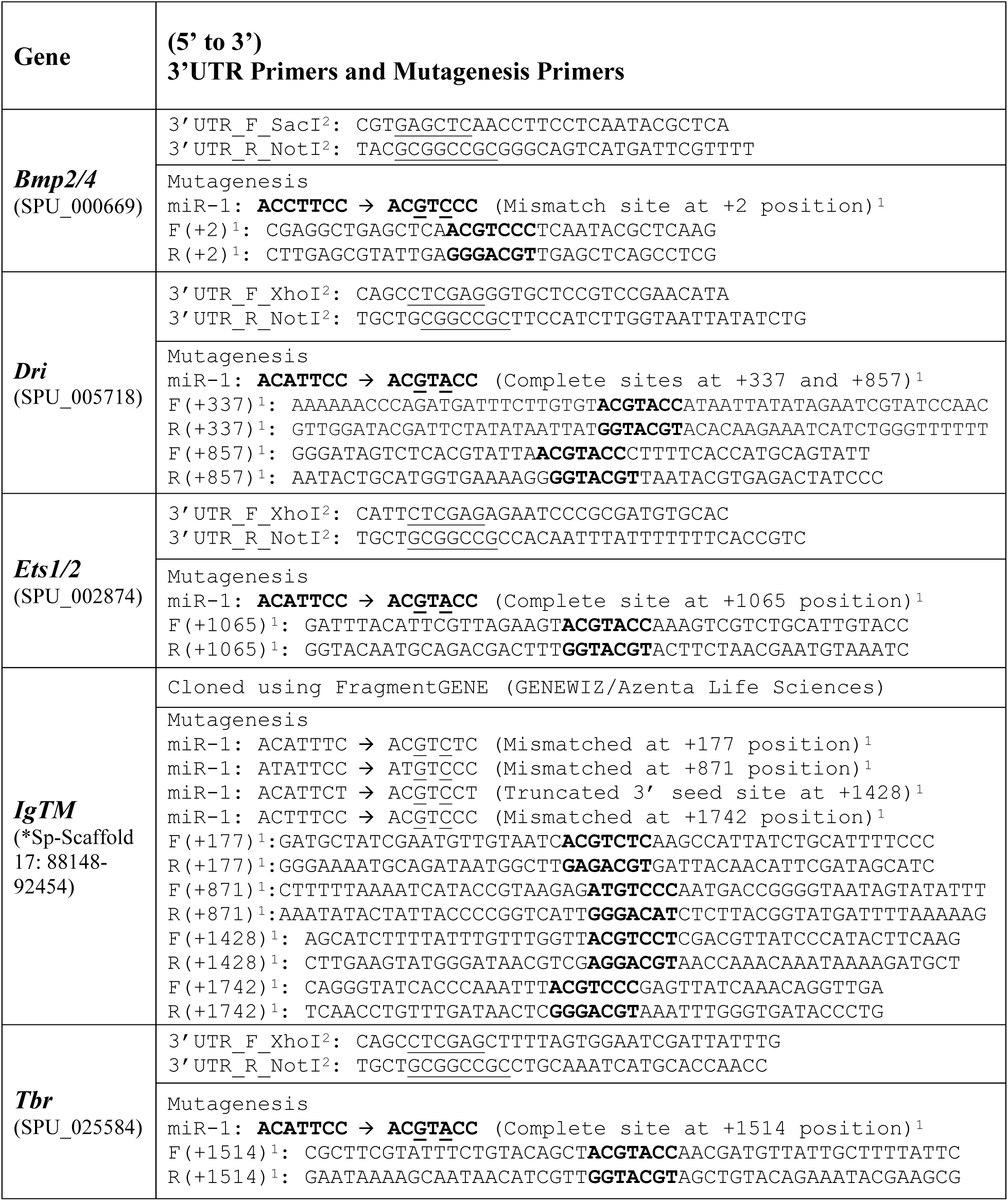
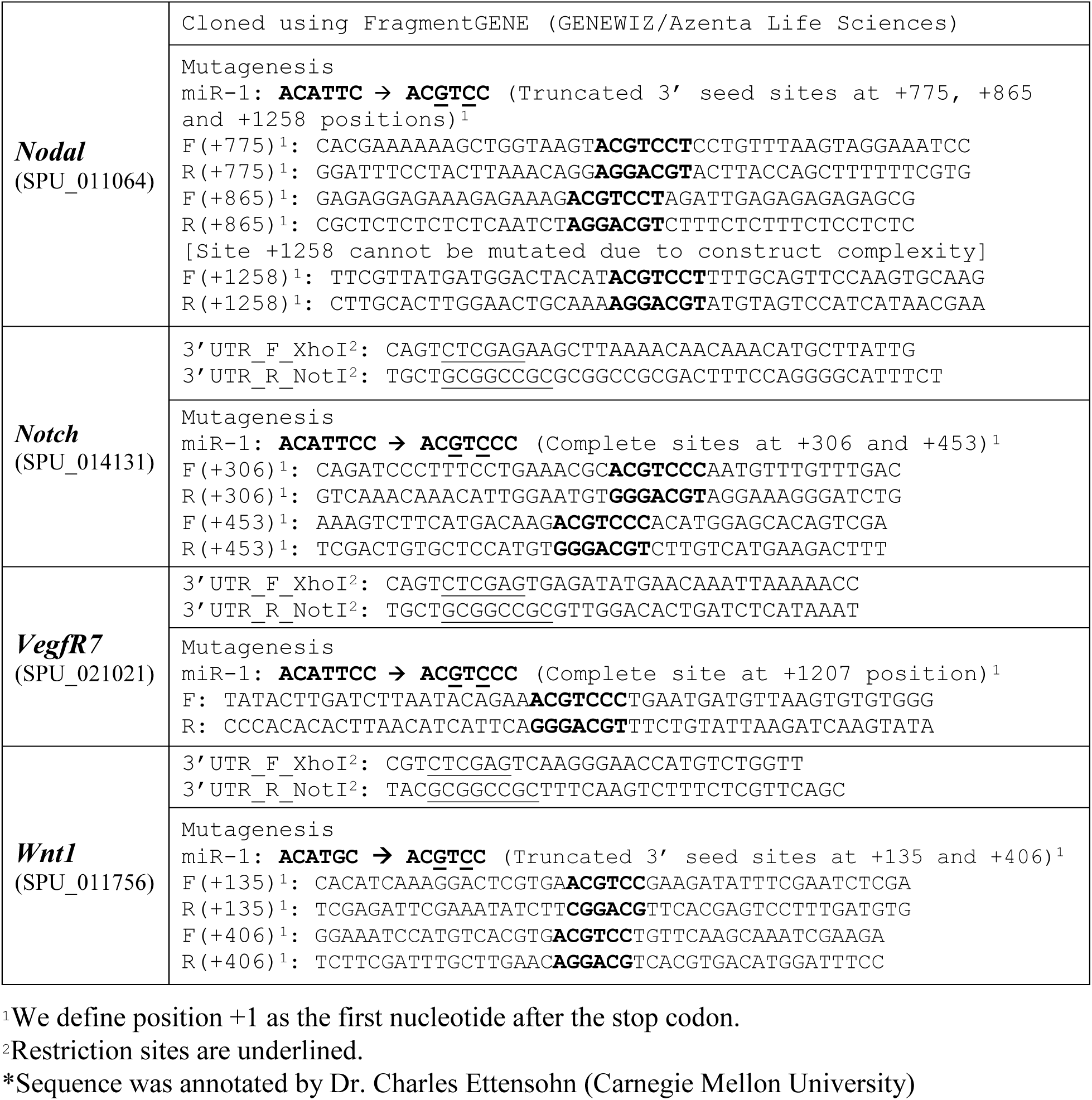
Primers for 3’UTR cloning and modifications of miRNA seed sequences of target genes for site-directed mutagenesis

**Table S2.**
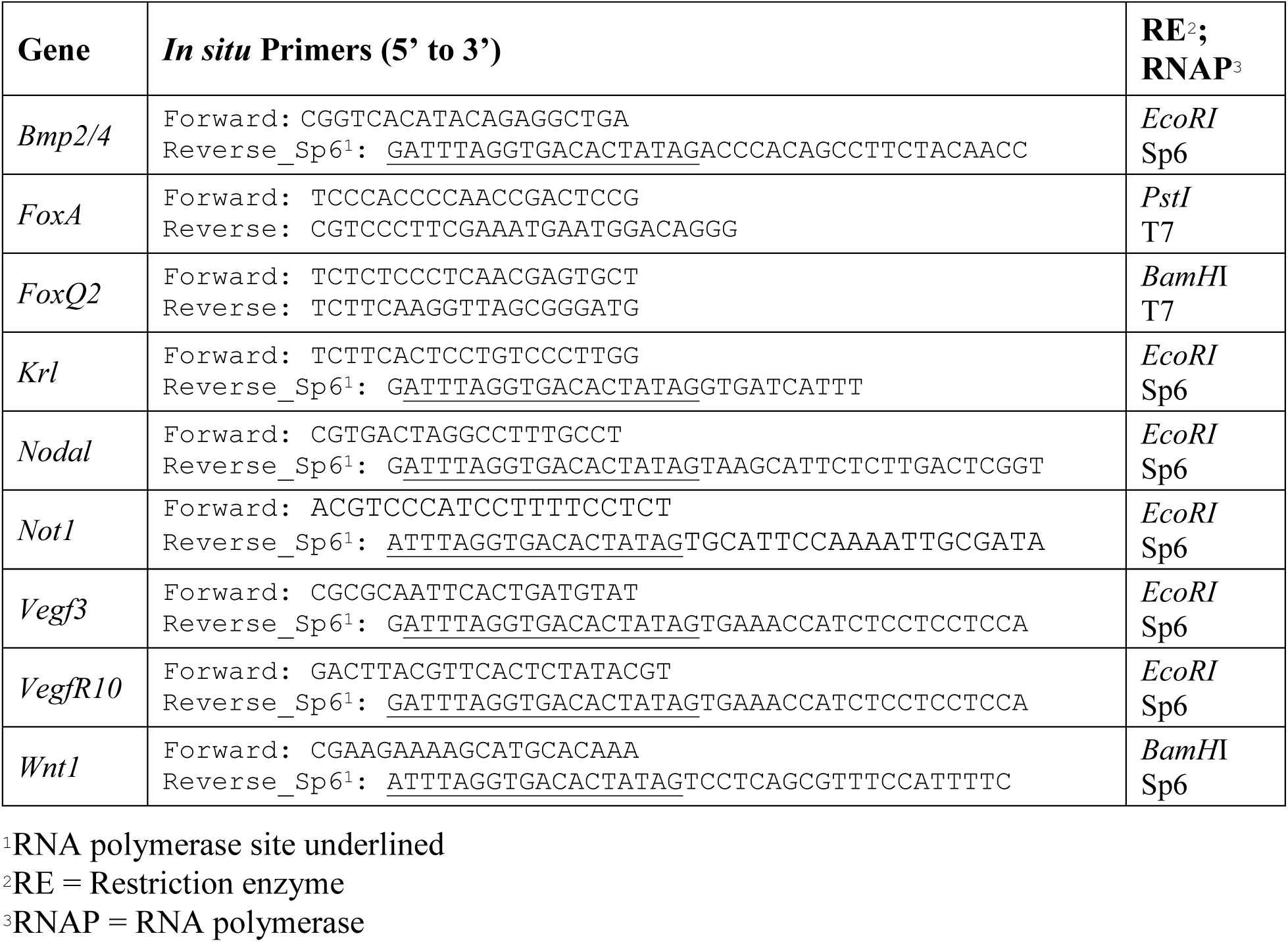
Primers and enzymes used for *in situ* hybridization

**Table S3.**
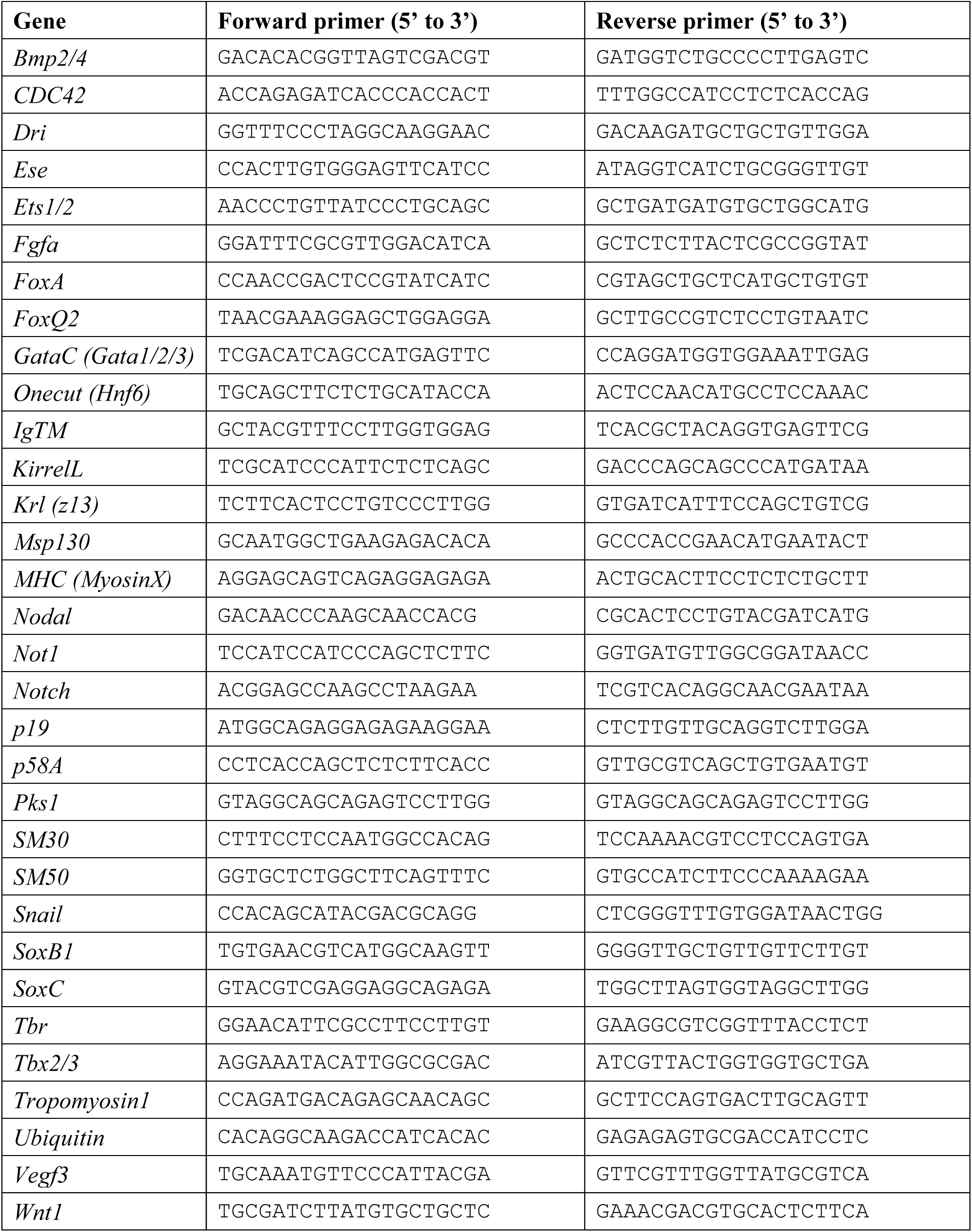
Relative real-time, quantitative PCR Primers

